# Cholesterol-containing liposomes decorated with Au nanoparticles as minimal tunable fusion machinery

**DOI:** 10.1101/2022.10.28.514049

**Authors:** Ester Canepa, Davide Bochicchio, Paulo Henrique Jacob Silva, Francesco Stellacci, Silvia Dante, Giulia Rossi, Annalisa Relini

## Abstract

Membrane fusion is essential for the basal functionality of eukaryotic cells. In physiological conditions, fusion events are regulated by a wide range of specialized proteins, as well as by a finely tuned local lipid composition and ionic environment. SNARE proteins, for example, provide the mechanical energy necessary to achieve vesicle fusion in neuromediator release, and their action is assisted by other soluble proteins, membrane cholesterol, and calcium ions. Similar cooperative effects must be explored when considering synthetic approaches to achieve controlled and selective membrane fusion. Here we show that liposomes decorated with amphiphilic Au nanoparticles (AuLips) can act as minimal tunable fusion machinery. AuLips fusion is triggered by divalent ions, while the number of fusion events dramatically depends on, and can be finely tuned by, the liposome cholesterol content. Our results, obtained via a combination of experimental (Quartz-Crystal-Microbalance with Dissipation monitoring, Fluorescence assays, Small-Angle X-ray Scattering) and computational techniques (Molecular Dynamics with coarse-grained resolution), reveal new mechanistic details on the fusogenic activity of amphiphilic Au nanoparticles in synergy with membrane cholesterol, and demonstrate the ability of these synthetic nanomaterials to induce fusion regardless of the divalent ion used (Ca^2+^ or Mg^2+^). This evidence provides a novel contribution to the development of new artificial fusogenic agents for next-generation biomedical applications that require tight control of the rate of fusion events (e.g., targeted drug delivery).

## INTRODUCTION

Membrane fusion is essential for the basal functionality of eukaryotic cells. Synaptic transmission^1,2^, membrane trafficking^3–5^, and subcellular cargo transport^5,6^ are some physiological events involving rapid and selective membrane fusion. In pathological conditions, membrane fusion is implicated in cancer progression^7^ and viral infection^8,9^, and defects in its machinery may result in severe biological damage, including disorders in neuronal synapses^10^. In recent years, controlled fusion *in vitro* became important from a biotechnology standpoint, as well: many efforts are being devoted to optimizing the fusion process between synthetic liposomes loaded with therapeutic agents and secreted extracellular vesicles and thus building non-immunogenic drug delivery platforms^11–13^.

Whether involving liposomes, secreted vesicles, or cells, the fusion process evolves through a few basic steps in all membrane types. First, fusion requires that the two membranes approach each other at < 5 nm distance. Then, fusion is initially triggered by the appearance of a bilayer stalk between the outer leaflets of the two interacting membranes^14–17^. The fusion stalk allows for mixing outer leaflet lipids and represents the first intermediate structure encountered in the fusion pathway^15,18–20^. The stalk then expands into a hemifusion diaphragm that eventually breaks to form the fusion pore that allows the mixing of inner-leaflet lipids and aqueous contents^15,16,18,21^. The fusion kinetics is ruled by energy barriers, which must be overcome to complete the fusion process^17,19,20^.

Although membrane fusion is universal, its mechanism is tightly regulated by distinct repertoires of biological components that vary from system to system. This is the case, for instance, of SNARE (Soluble N-ethylmaleimide-sensitive factor Attachment protein REceptors) proteins – peripheral membrane components involved in the initiation of membrane fusion reactions in secretory pathways such as exocytosis and synaptic transmission^21–23^. SNAREs are responsible for stabilizing the first stages of membrane juxtaposition, leading to inter-membrane distances of 4-6 nm. Other protein components, membrane-tethered (e.g., synaptotagmin-1) and soluble (e.g., complexin), can form stable complexes with SNAREs and favor the advancement of the fusion process through its different stages^24,25^. The fusogenic action of these protein complexes is, in turn, influenced by two other players: ions and lipids. Ions play essential regulatory roles in many membrane fusion processes in cells and natural or synthetic vesicles. Calcium ions, for instance, are crucial in promoting synaptic vesicle fusion in synergy with SNARE assemblies^24,26^, altering both protein-protein affinities and protein conformations^24^. As for lipids, cholesterol plays a crucial role in promoting SNARE-mediated fusion. It has been shown to favor vesicle exocytosis in neuronal, endocrine, and neuroendocrine cells^27^, and its removal significantly impairs secretory content release^27^. The effect of cholesterol has different origins and is, in part, still debated. On the one hand, cholesterol fusogenicity is due to specific interactions with the proteins at the fusion site, as cholesterol promotes clustering of SNAREs^27^; on the other hand, owing to its intrinsic negative curvature, cholesterol could lower dehydration barriers^28^. The negative curvature of cholesterol may also have a role in stabilizing the metastable stalk state, even though there is no evidence of an accumulation of cholesterol in the stalk region^20^, as well as in shortening the lifetime of the hemifusion intermediate state, leading to fast fusion pore opening^29^ and reduced pore flickering^30^.

Only recently, it has been revealed that artificial agents can also act as effective biomimetic fusogens. This is the case of complementary DNA strands, or peptide sequences^31,32^, as well as engineered nanoparticles (NPs) with solid metal-oxide or metal core ^33–35^. The latter include sub-5 nm gold NPs (AuNPs) protected by an amphiphilic thiol monolayer comprising the negatively charged 11-mercapto-1-undecanesulfonate (MUS) and the apolar 1-octanethiol (OT)^36,37^. The amphiphilic MUS:OT surface functionalization is extremely promising for biomedical purposes, as it allows for spontaneous NP penetration into biomimetic lipid bilayers^38–44^ and mammalian cells^45–50^ in a non-destructive way and the possibility of conjugating drug cargoes for intracellular delivery^36,50,51^. More interestingly, MUS:OT AuNPs recently demonstrated to enable Ca^2+^-triggered fusion of pure phosphatidylcholines (PC) membranes^33^ – a peculiar effect that does not occur when neutral zwitterionic membranes come into contact with calcium alone^52,53^.

Here we show, by a combined experimental and computational approach, that the fusogenic activity of these AuNPs can be dramatically enhanced, and thus finely tuned, by the combined action of divalent ions and membrane cholesterol. We thus propose that cholesterol-containing liposomes decorated with bilayer-embedded MUS:OT AuNPs (AuLips) may be regarded as minimal hybrid fusion machinery. In the following sections, we present our data obtained through Quartz Crystal Microbalance with Dissipation monitoring (QCM-D), fluorescence-based membrane fusion assays, and Small-Angle X-ray Scattering (SAXS), showing that membrane cholesterol dramatically enhances the fusogenic activity of MUS:OT AuNPs. Furthermore, we use Molecular Dynamics (MD) simulations to gain a molecular-level understanding of the contribution of cholesterol to the membrane fusion process, both in terms of kinetics and thermodynamics. Interestingly, our results also reveal the ability of these synthetic NPs to induce fusion regardless of the divalent ion used (Ca^2+^ or Mg^2+^). We believe these results represent a fundamental step towards understanding how engineered NPs can be rationally designed to achieve finely controlled fusion events in artificial and living membranes for next-generation biomedical applications.

## RESULTS

### AuNPs and cholesterol-containing liposomes

We synthesized and used monodisperse AuNPs with an average core diameter of 2.4 nm and protected by an amphiphilic monolayer of MUS and OT ligands mixed in a 2:1 molar ratio (Figures S1,S2 and Table S1)S1^43^. The structure of these MUS:OT AuNPs is shown in Figure S3, together with the AuNP coarse-grained model used in our Molecular Dynamics simulations. To mechanistically investigate the peculiar fusogenic activity of MUS:OT AuNPs, we used 100 nm extruded fluid-state liposomes made of 1,2-dioleoyl-*sn*-glycero-3-phosphocholine (DOPC) and biologically relevant percentages of cholesterol (0÷40 mol %)^43,54^ (Figure S4).

### QCM-D, fluorescence, and SAXS experiments

Membrane cholesterol has recently been shown to hinder the passive penetration of sub-5 nm MUS:OT AuNPs within fluid PC bilayers^43^. In particular, as the cholesterol content increases, the number of amphiphilic NPs that manage to enter the lipid bilayer is drastically reduced, leading to potential implications for their fusogenic activity in membrane fusion studies. In order to separate the contribution of membrane cholesterol in modulating NP-promoted membrane fusion from effects due to the number of membrane-embedded NPs, we quantified by QCM-D the membrane incorporation of 2.4 nm MUS:OT AuNPs in our experimental conditions. Based on previous studies by our group, we relied on supported vesicle layers (SVLs) deposited on the surface of a gold-coated QCM sensor^43^. On gold, vesicles do not merge and form viscoelastic vesicle layers that are ideal for quantifying mass variations due to the vesicles’ ability to incorporate NPs passively. The QCM-D monitoring over time of the passive NP uptake into cholesterol-enriched DOPC vesicles in 2.5 mM Trizma® base and 50 mM NaCl (pH 7.4, 25° C) is reported in Figure 1. Figure 1a shows a representative example of our QCM-D experiments recorded at an intermediate cholesterol percentage (20 mol %). At all cholesterol concentrations, NPs were inserted into the QCM chamber after rinsing off the excess vesicles left by SVL deposition. As evidenced by Figure 1a, NP addition induced an immediate and detectable reduction in the SVL normalized resonance frequency (Δf_norm_), corresponding to an increase in the mass deposited on the sensor surface. This trend is consistent with the known ability of sub-5 nm 2:1 MUS:OT AuNPs to passively enter the fluid bilayer of PC vesicles^40–42,44^. Here, the spontaneous NP uptake is also confirmed by the remarkable stability of the SVL–NP complex after the final rinsing of the samples (Figure 1a). The SVL mass variations monitored by QCM-D (Δm_SVL_ %) are shown in Figure 1b. These data confirm the tendency of membrane cholesterol to reduce passive NP incorporation into unsaturated phospholipid vesicles and are in excellent agreement with previous results obtained in different salt buffers (i.e., 137 mM NaCl)^43^.

**Figure 1.**
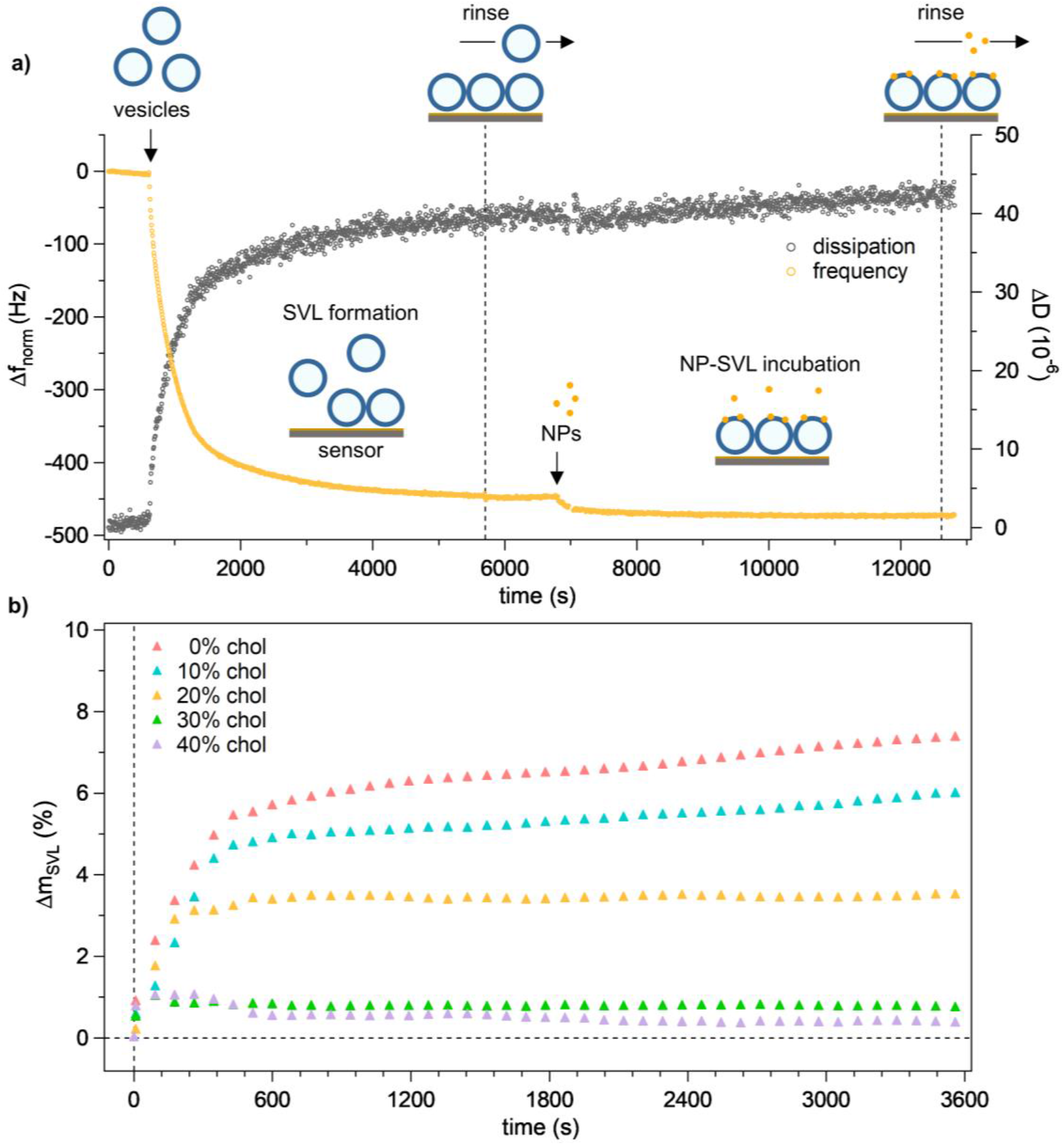
Dissipative QCM investigation (pH 7.4, 25°C) to monitor the passive uptake of amphiphilic NPs into supported vesicle layers (SVLs) as membrane cholesterol content (mol %) increases. **a)** Normalized frequency (Δf_norm_) and dissipation (ΔD) traces (3^rd^ overtone) of a representative experiment before and after NP addition (20 mol % chol; 7 s sampling interval). Vesicles were inserted into the QCM chamber after 600 s of thermal equilibration, while NPs were injected after rinsing the preformed SVL (system cartoon not to scale). The two reductions in sensor oscillation frequency at 600 s and 6800 s correspond to the increase in mass deposited on the sensor surface due to SVL formation and NP uptake, respectively. **b)** Monitoring over time of the percentage change in SVL mass (Δm_SVL_) during NP incorporation (NPs added at t=0 s). Δm_SVL_ (%) results – smoothed in (b) by a 20-point binomial filter – were obtained from Δf_norm_ data. See Supplementary Information for full details on sample preparation, experimental setup, and data analysis.

As shown in Figure 1b, depending on its cholesterol content, the vesicle lipid bilayer exhibits a very different ability to accommodate amphiphilic NPs passively. In particular, in the absence of cholesterol or at molar percentages below 20 %, spontaneous NP incorporation shows similar trends in the first few minutes before asymptotically reaching distinct maxima. At higher cholesterol percentages, however, only a few NPs can penetrate the vesicle lipid bilayer, with plateau values drastically lower than those obtained in more fluid membranes. Based on this evidence, we subsequently conducted membrane fusion studies with a NP-vesicle incubation time limited to 5 minutes, for which the difference between NP uptake in the absence of cholesterol and at maximum cholesterol content was reduced to ∼4:1. We probed membrane fusion by conventional assays based on fluorescence resonance energy transfer (FRET) and fluorescence dequenching (Figure 2a)^33,55–57^. These fluorometric methods were exploited to measure the mixing of vesicle lipids and aqueous contents after CaCl_2_ addition, respectively. As in previous membrane fusion studies^33^ and common neurophysiology experiments^58^, Ca^2+^ ions were added at an extravesicular concentration of 2 mM. This value is representative of the total calcium concentration in the extracellular milieu (∼10^−3^ M)^58,59^ and has been shown to promote significant mechanical stress and structural changes in the bilayer of pure zwitterionic PCs^52^.

**Figure 2.**
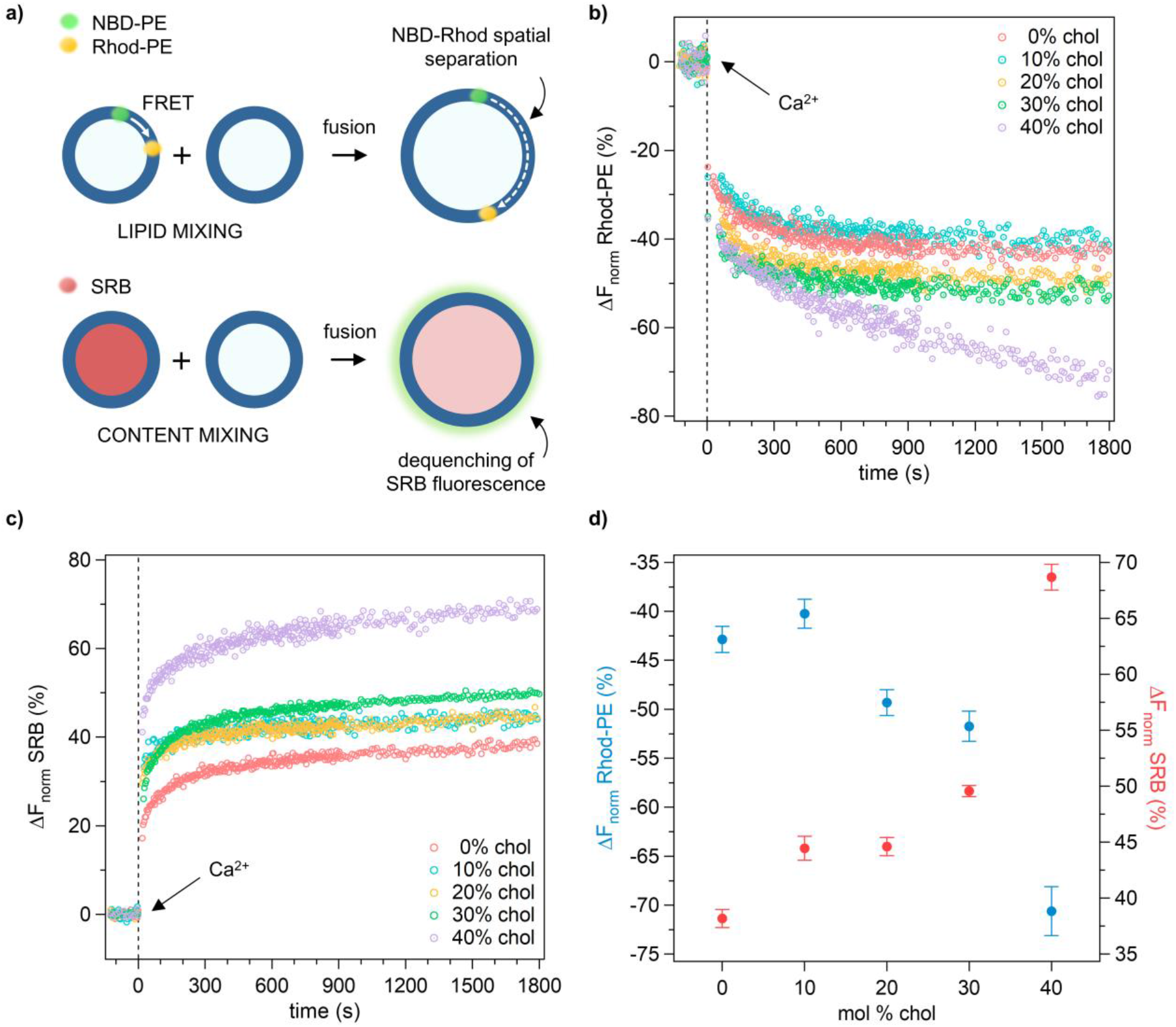
Real-time fluorescence assays to probe vesicle fusion in the presence of membrane-embedded NPs and increasing membrane cholesterol content (pH 7.4, 25°C). **a)** Schematic drawing illustrating the lipid and content-mixing assays used in this study; both assays involve the interaction between fluorescently labeled and unlabeled vesicles. Lipid mixing is assessed by measuring the dilution-related reduction in proximity-dependent FRET signal between the donor (NBD-PE) and the acceptor (Rhod-PE) fluorescent lipids. Content mixing, on the other hand, is tested by detecting dilution-related dequenching of a fluorescence dye (SRB) loaded within the vesicle lumen. **b)** Normalized reduction (%) of Rhod-PE emission due to dilution of fluorescent lipids during mixing of vesicles’ inner leaflets. **c)** Normalized increase (%) of SRB emission due to dilution of the self-quenched dye during mixing of vesicles’ contents. In (b) and (c), Ca^2+^ ions (2 mM) were added after 5 min of NP-vesicle incubation; temporal changes in the fluorescence traces were then monitored for 1800 s (4 s sampling interval before 900 s, 10 s after). Importantly, no change in FRET signal (b) and SRB emission (c) was recorded before Ca^2+^ addition. **d)** Lipid and content-mixing results (mean ± std. error) averaged over the last 5 min of recording in (b) and (c). See Supplementary Information for full details on sample preparation, experimental setup, and data analysis.

In lipid mixing assays, inserting two FRET probes into the vesicle bilayer allowed us to monitor temporal changes in membrane composition during the interaction between labeled and unlabeled vesicles (Figure S5a)^60,61^. At first, using the donor lipid probe NBD-PE and the acceptor lipid probe Rhod-PE, we verified that Ca^2+^ alone (in the absence of NPs) does not promote vesicle lipid mixing (Figure S6). On the contrary, Ca^2+^ added after 5 min of NP-vesicle incubation resulted in an immediate reduction of the proximity-dependent FRET signal at all cholesterol concentrations. This is revealed by the clear suppression of the acceptor probe’s emission reported in Figure 2b and is evidence that the membrane lipids of interacting vesicles have been diluted. Notably, the FRET decrease shown in Figure 2b became increasingly more pronounced as the membrane cholesterol content increased in the molar range 10÷40 %. Representative emission spectra acquired during lipid mixing assays are shown in Figure S7. Since lipid mixing could be compatible with incomplete fusion – i.e., with the formation of stalked states in which only the outer leaflets of the vesicle membranes mix their lipids – we treated our vesicle system with sodium dithionite, a specific quencher of NBD fluorescence emission^62^. When we added the fresh NBD quencher to the external buffer, it did not induce relevant changes in the FRET signal even at high concentrations (> 50 mM), indicating that the NBD labeling of the lipid bilayer mainly involved the inner leaflet of our PC vesicles (Figure S8). This observation is clear evidence that amphiphilic NPs can promote, in the presence of Ca^2+^ ions, intervesicle mixing of inner-leaflet lipids.

We further investigated NP-mediated vesicle fusion by content mixing assays. Here, unlabeled vesicles were let to interact with vesicles loaded with the fluorescent dye Sulforhodamine B (SRB) (Figure S5b)^33,56,57,63,64^. By encapsulating a self-quenching concentration (50 mM) of this small water-soluble fluorophore within the aqueous vesicle content, we monitored any enhancement in sample fluorescence caused by probe dilution in real-time. As expected, calcium alone proved ineffective in promoting changes in sample fluorescence without NP-vesicle incubation (Figure S9). However, as shown in Figure 2c, after 5 min of NP-vesicle incubation, the addition of Ca^2+^ induced a sudden rise in SRB emission due to fluorescence dequenching. A typical content mixing trace before data normalization is reported in Figure S10. Consistent with the trend in lipid mixing highlighted in Figure 2b, the increase in SRB fluorescence became more pronounced at higher cholesterol contents. These results suggest that Ca^2+^-triggered redistribution of inner-leaflet vesicle lipids enabled by membrane-embedded NPs (Figure 2b) is accompanied by the opening of fusion pores and subsequent mixing of aqueous contents between labeled and unlabeled vesicles. As better detailed in the Supplementary Information, we performed control experiments to relate this evidence to complete membrane fusion directly and exclude that increased SRB emission was mainly due to vesicle leakage (Figure S5c). Although leakage assays indicate that there is some dilution of the SRB probe with the external medium due to partial destabilization of the vesicle bilayer (Figure S11), this is not the main cause of SRB dequenching in our content mixing assays.

The overall results of our membrane fusion assays are summarized in Figure 2d. These data demonstrate that the ability of MUS:OT AuNPs to enable Ca^2+^-triggered fusion of pure PC membranes is retained at biologically relevant concentrations of membrane cholesterol (0÷40 mol %). Most importantly, our investigations show that an increase in cholesterol content, despite strongly stiffening the fluid bilayer of the PC membrane^43^ and reducing the passive uptake of amphiphilic NPs (Figure 1b), significantly enhances their fusogenic potential. We further investigated this evidence by probing how the amount of Ca^2+^ ions affects the NP fusogenic activity. Specifically, employing lipid mixing assays, we tested decreasing concentrations of Ca^2+^ ions from 2 mM up to 100 µM on vesicles incubated with NPs. This range is biologically relevant as it is representative of the strong concentration gradient generated by the entry of Ca^2+^ ions into cells through specific ion channels in the plasma membrane^65^. Indeed, in the vicinity of entry sites, the intracellular Ca^2+^ level is ∼10^−4^÷10^−5^ M, while it drops to ∼10^−7^ M over a few hundred Å within the cytosol^59,65^. As shown in Figures 3a-c, our investigation revealed a pronounced Ca^2+^-ions concentration dependence on the ability of the NP to promote vesicle fusion. In particular, the fusogenic activity of NPs intensifies significantly as the concentration of extravesicular Ca^2+^ increases in the 100 µM÷2 mM range (Figure 3a,b) – a tendency shared by both cholesterol-free vesicles and those containing increasing amounts of membrane cholesterol (Figure 3c). Taken together, this Ca^2+^-dependence of fusion results suggests that the fusogenic activity of MUS:OT AuNPs should be maximized near the outer leaflet of plasma membranes containing high cholesterol levels, such as those of neuronal cells^66^.

**Figure 3.**
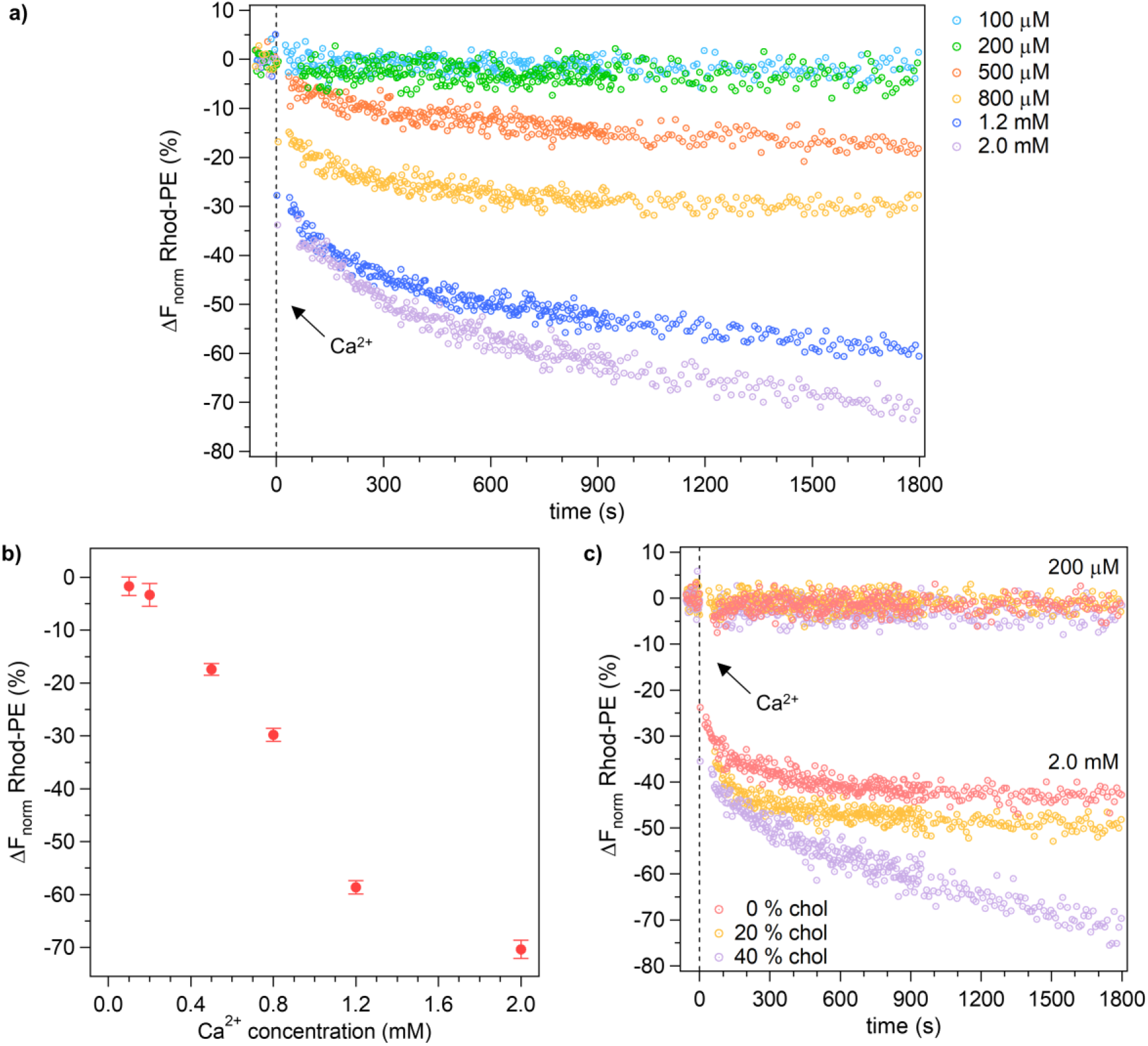
The extravesicular Ca2+ concentration tightly regulates NP-promoted vesicle fusion enhanced by membrane cholesterol. **a)** Normalized reduction (%) of Rhod-PE emission recorded after addition of increasing concentrations of Ca^2+^ ions (0.1÷2 mM) to high-cholesterol vesicles (40 mol % chol) pre-incubated with NPs (pH 7.4, 25°C). **b)** Lipid mixing results (mean ± std. error) averaged over the last 5 min of the traces shown in (a). **c)** Normalized reduction (%) of Rhod-PE emission due to lipid mixing induced by low (200 µM) and high (2.0 mM) levels of Ca^2+^ in vesicles pre-incubated with NPs and containing three representative concentrations of membrane cholesterol: none (0 mol %), medium (20 mol %) and high (40 mol %). In (a) and (c), calcium was added after 5 min of NP-vesicle incubation; temporal changes in real-time fluorescence traces were then monitored for 1800 s (4 s sampling interval before 900 s, 10 s after). For full details on sample preparation, experimental setup and data analysis, see Supplementary Information.

We further evaluated NP-promoted vesicle fusion by performing SAXS measurements. Experiments were conducted on 50 nm extruded DOPC vesicles (10 mg/mL) with increasing molar content of membrane cholesterol (0, 20, 30, and 40%). Each sample was measured before NP addition, after incubation with diluted NPs (∼1 NP/vesicle), and after administration of Ca^2+^ ions (2 mM). Results are shown in Figure 4 and Figures S12-13 (see also Table S2). Regardless of cholesterol content, NP addition induced an increase in the scattering signal in the low momentum transfer (*q*) region, thus indicating an increase in the size of the scattering objects^67^. Ca^2+^ administration, on the other hand, affected the scattering curves in a cholesterol-dependent manner: for cholesterol percentages up to 20 mol %, the scattering curves before and after Ca^2+^ addition were superimposable (Figure 4a and Figure S12); at higher cholesterol percentages (30 and 40 mol %), the scattering signal in the low-*q* region decreased significantly after the addition of divalent ions. To quantify these effects, the experimental curves of lipid vesicles were fitted using the usual spherical model with 3 shells – i.e. the regions of the lipid bilayer (see Figure S13)^68–70^.

**Figure 4.**
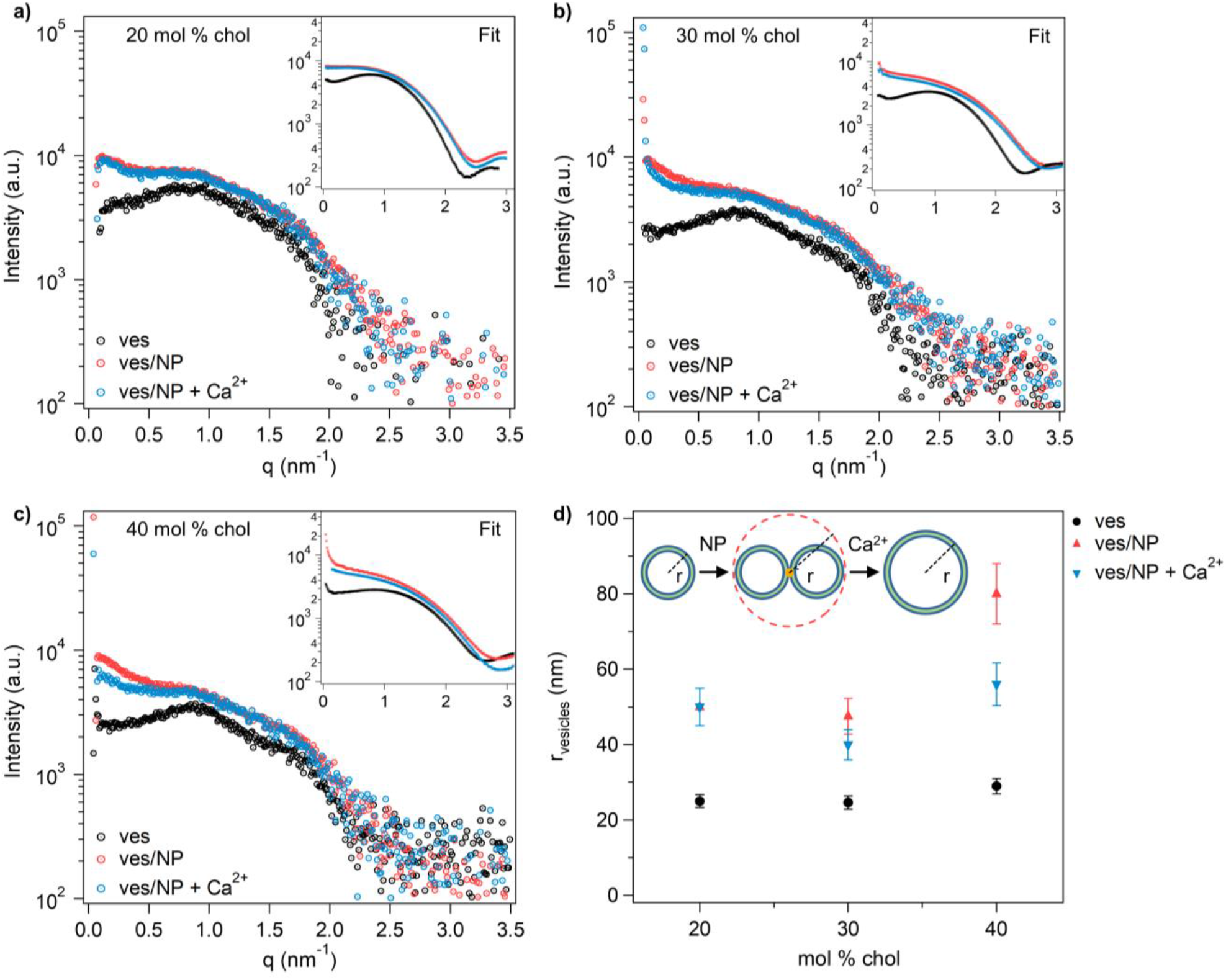
SAXS curves obtained from 50 nm extruded DOPC vesicles with cholesterol molar content of **a**) 20%, **b**) 30% and **c**) 40% (pH 7.4, 25°C). Pure vesicles (black circles) fitted with a spherical core-shell model returned a vesicle radius *r* of 25 nm (**d**); after NP incubation, the scattering curves (red circles) are characterized by the higher intensity at low-*q*, while after Ca^2+^ administration the scattering intensity in the same region (light blue circles) decreased as a function of cholesterol content. The simulated curves are displayed in the insets. The vesicle sizes resulting from the fitting procedure are reported in panel (**d**). A sketch of the suggested fusion mechanism is reported in the same panel.

The results of the fitting procedures are shown in the insets of Figure 4a-c. All details about the fitting procedures are in the Supplementary Information. The vesicle size returned by the fits as a function of cholesterol percentage is shown in Figure 4d. As expected, the radius (*r*) of the vesicle core in the pure vesicle samples is in perfect agreement with the size of the extrusion pores, i.e. ∼25 nm. As anticipated earlier, after incubation with NPs, the size of the vesicles increased from the initial radius of 25 nm to a maximum value of 80 nm at 40 mol % chol. This size increase is consistent with the formation of docked vesicles through membrane-adsorbed NPs, as depicted in the sketch in Figure 4d. The aggregation of lipid vesicles promoted by MUS:OT AuNPs is a known phenomenon already observed in previous experimental evidence^33,44^. The polydispersity of the vesicle size after incubation with NPs is also increased, with respect to pure vesicles (Figure 4d). After the addition of Ca^2+^ ions, only for the highest cholesterol contents (30 and 40 mol %), the vesicle core size decreased. This effect is interpreted as the fusion of the docked vesicles, giving rise to a single smaller vesicle. We emphasize that we used a very low NP/vesicle ratio (∼1) to avoid the formation of large NP aggregates in the solution that would form after the addition of Ca^2+^ ions, as previously observed from DLS measurements (Figure S2). Additionally, the accessible instrumental *q*-range allows the detection of objects as large as 100 nm, compatible with aggregates of up to 2–3 vesicles. Therefore, the low NP/vesicle ratio also hinders the formation – mediated by numerous NPs – of large vesicle aggregates with a size beyond the instrumental accessibility.

In addition to fluorescence and SAXS, membrane fusion was further verified using the QCM-D setup discussed above (Figure 1a), which offers the advantage of monitoring vesicle fusion without NPs in solution. Once the SVL was deposited and allowed to interact with the NPs, we thoroughly rinsed the sample to remove excess NPs that had not penetrated the vesicle bilayer. Next, we inserted into the QCM chamber Ca^2+^ ions (2 mM) mixed with fresh vesicles (Figure 5a and Figure S14). Only 100 nm extruded vesicles were employed for this QCMD study. As shown in Figure 5a, adding vesicles and Ca^2+^ ions induced a significant decrease in the sensor’s normalized resonance frequency, consistent with the fusion of SVL’s vesicles with those in solution and the formation of larger (and thus heavier) fused vesicles on the sensor surface.

**Figure 5.**
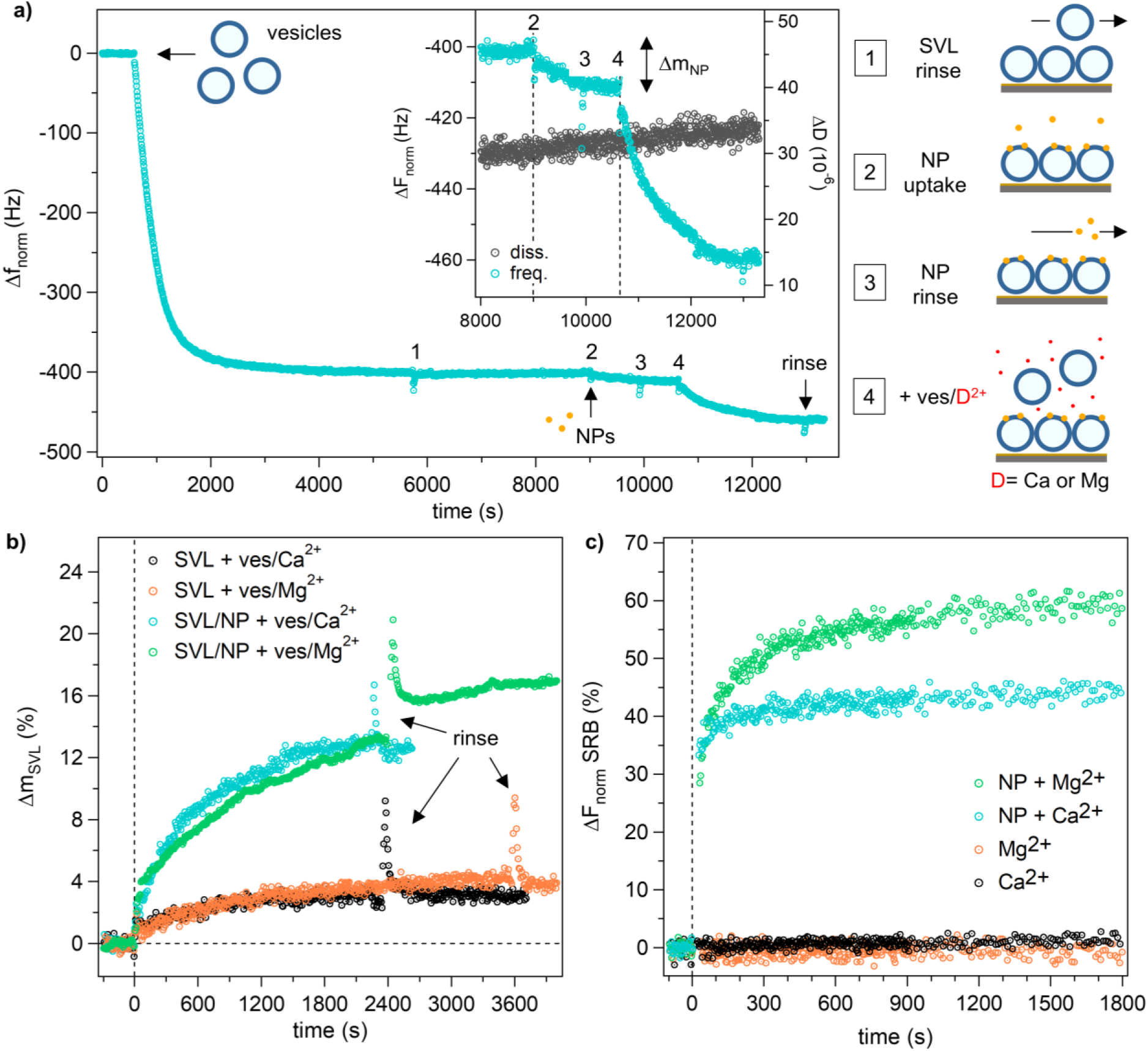
Vesicle fusion is promoted by membrane-embedded NPs in the presence of both Ca^2+^ and Mg^2+^ ions (pH 7.4, 25°C). **a)** Representative QCM-D membrane fusion experiment conducted on an SVL containing 10 mol % chol (7 s sampling interval). After copious rinsing of excess NPs in solution, additional vesicles (0.05 mg/mL, 10 mol % chol) incubated with Ca^2+^ 2 mM were added to the QCM-D chamber (cartoons not to scale). The inset shows the enlarged experimental trace between NP addition and final post-fusion SVL rinse. The two reductions in the sensor oscillation frequency (steps 2 and 4) are due, respectively, to NP uptake by the SVL (Δm_NP_) and to the divalent ion-induced fusion between the supported vesicles containing NPs and those freshly inserted in solution. **b)** Monitoring over time of the percentage change in the mass of the SVL-NP complex containing 10 mol % chol (Δm_SVL_) after addition at t=0 s of fresh vesicles incubated with Ca^2+^ or Mg^2+^ ions (7 s sampling interval). In the absence of NPs, minimal mass perturbations were recorded after vesicles/ions addition. **c)** Confirmation by content mixing assays of the fusogenic effect of Mg^2+^ ions (2 mM) on PC vesicles (10 mol % chol) after 5 min NP incubation (4s sampling interval before 900 s, 10 s after). For full details on sample preparation, experimental setup, and data analysis, see Supplementary Information.

Finally, the same QCM-D system was used to probe whether the fusogenic ability of MUS:OT AuNPs exhibits any specificity towards divalent Ca^2+^ or Mg^2+^ ions. Indeed, the latter are generally known to exert a much more limited fusogenic activity on phospholipid vesicles with different compositions^53,71–73^. Surprisingly, QCM-D experiments repeated by replacing Ca^2+^ with Mg^2+^ provided comparable results, revealing that magnesium ions can also trigger the fusion of PC-based vesicles when NPs are present (Figure 5b). The emerging ability of MUS:OT AuNPs to induce fusion of PC vesicles even in the presence of Mg^2+^ ions was confirmed by the content mixing experiments shown in Figure 5c. Further analyses of NP-promoted membrane fusion are discussed in Figure S15 of the Supplementary Information.

### In silico experiments

To further investigate the role of cholesterol in favoring the NP-induced membrane fusion and add molecular details to the whole picture, we moved to molecular dynamics (MD) investigations exploiting sub-molecular resolution coarse-grained (CG) models. As done in a previous study,^43^ we modeled the lipid bilayers and the MUS:OT AuNPs (Figure S3) based on the Martini CG force field.^74^ In particular, our computational investigations focused on the first step of the fusion process: the formation of the stalk.

We prepared and equilibrated two alternative systems composed of two parallel DOPC (**S1**) or DOPC + 30% mol cholesterol (**S2**) lipid bilayers, including a single MUS:OT NP partially snorkeled into the bottom bilayer (Figure 6a). In both systems, we set the distance between the two bilayers to ∼ 2 nm, allowing the NP to interact spontaneously with the top bilayer through its charged terminals of the MUS ligands (Figure 6a, *State A*). Then, we performed 8 independent, 4 µs long unbiased MD simulations in the NPT ensemble for each system using semi-isotropic pressure scaling. The temperature was set to 370 K to accelerate the stalk formation, which is an activated process. Indeed, despite using a CG model, the free energy barrier of stalk formation remained too high to be overcome at 310 K in reasonable simulation times. However, aiming to compare the **S1** and **S2** systems and not to obtain absolute quantities, it was sufficient to use the same temperature in both cases.

**Figure 6.**
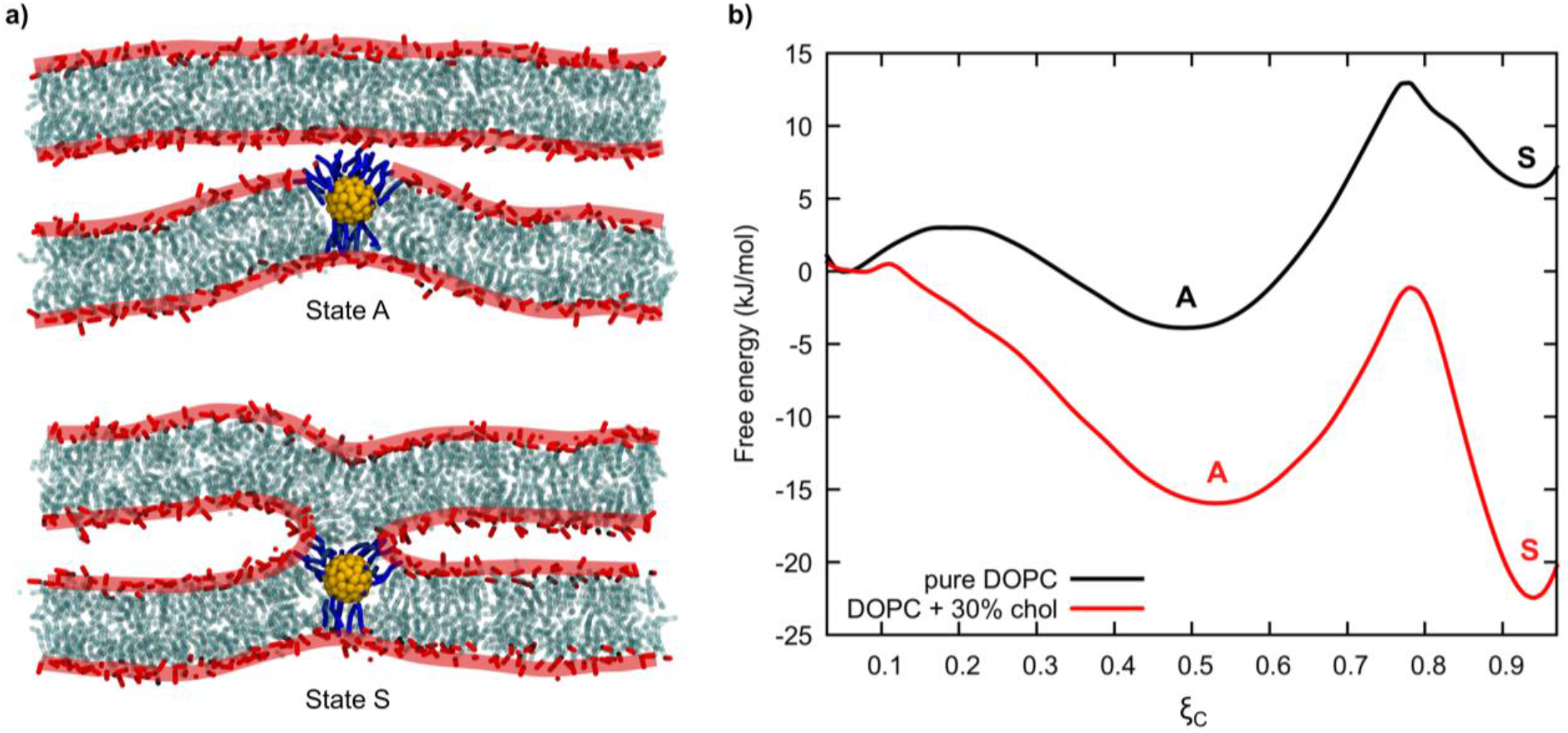
Stalk formation investigated with MD simulations employing CG models. a) Snapshots from a representative MD simulation of the **S2** system, showing the initial configuration (NP adsorbed onto the bilayer above, state A) and the final configuration (after 4 µs) in which the stalk is formed (state S). For clarity, we show only DOPC polar heads (in red), DOPC hydrophobic tails (in cyan), the Au core of the NP (in yellow), and the MUS ligands (in blue). The boundaries of the two membranes are highlighted in red to stress the difference between the two configurations. b) Free energy profiles associated with the stalk formation process for the **S1** and **S2** systems. Minima corresponding to *state A* and *state S* are indicated with the respective letters.

In **S2** systems, we observed the formation of a stalk configuration (*State S*) in the microsecond time scale (Figure 6a). The stalk always formed above the NP, following the formation of hydrophobic contacts between the lipid tails of the top bilayer and the hydrophobic beads of the NP ligands. Once formed, the stalk in **S2** simulations always remained stable. On the contrary, the formation of a stalk was more sporadic in the **S1** system and, when formed, *state S* showed a short-lived metastable character (lifetime of a few nanoseconds), with the system spontaneously coming back to *state A*.

To be more quantitative, we developed a collective variable, ξ_C_, to analyze the stalk formation process. Inspired by the work of Hub and coworkers.^20^, we defined a cylinder placed at a fixed distance along the membrane normal above the NP and divided it into eight slices, as shown in Figure S16. ξ_C_ represents the average filling of the cylinder with hydrophobic lipid tail beads (see Supplementary Information for further details on the definition of ξ_C_ and the choice of the relevant parameters). Here we stress that, after tuning the parameters defining ξ_C_, we obtained a collective variable able to distinguish *State A* and *State S* in both **S1** and **S2** systems (Figure S17). In particular, ξ_C_ close to 1 means that the cylinder is full of lipid tails and thus that there is a well-formed stalk above the NP, while ξ_C_ close to 0 means that the cylinder is empty and thus that there is not a stalk.

Furthermore, to increase the statistics on the stalk configuration lifetime, we ran for both **S1** and **S2** systems a new set of 8 independent 4 µs long unbiased MD simulations starting from an already formed stalk (*state S*). In Table 1 we report the results on the stalk formation time and the stalk lifetime obtained from all the unbiased simulations. These results confirm our qualitative observations: the stalk is stable in the presence of cholesterol (**S2**), while its lifetime is really short in the case of pure DOPC bilayers (**S1**). Here we remind that we are using both CG models and a temperature higher than in the experiments, and thus all these computationally derived times are underestimated with respect to the experimental ones. However, interestingly, the stalk forms on the same timescale (µs) in both systems, indicating that cholesterol has a minor effect on the kinetics of stalk formation while having a major one on the thermodynamics (stalk stability).

**Table 1.** Average stalk formation time and stalk configuration lifetime, obtained from a total of 64 µs of unbiased MD at 370 K for each system (**S1** and **S2**). Standard errors of the mean are reported. We could not evaluate the average stalk lifetime in the **S2** system since there the stalk never disappeared once formed.

|  | Average stalk formation time<br>(ns) | Average stalk lifetime<br>(ns) |
| --- | --- | --- |
| DOPC (**S1** system) | 2250 (± 350) | 75 (± 15) |
| DOPC + 30% mol chol (**S2** system) | 1200 (± 350) | / |

Performing a simple Boltzmann inversion on the ξ_C_ data, we estimated the free energy profile of stalk formation in the two cases (Figure 6b). The free energy plots, while only semi-quantitative since we could not sample enough to observe the stalk destruction in the system with cholesterol, summarize very well our computational results and help interpret the experiments. For both S1 and S2, we obtain a profile with three free energy minima. On the left, for ξ_C_ < 0.1, we have a minimum corresponding to well-separated bilayers, in which the NP interacts only with the bottom one in which it is immersed. We have a minimum corresponding to *State A* at ξ_C_ = 0.5 and a minimum corresponding to *State S* at ξ_C_ > 0.9. The first barrier is related to the dehydration barrier. Here, our membrane patch is large enough to allow for a local, NP-induced membrane deformation that bring the NP to have short-range (< 1 nm) interactions with the facing membrane. The second barrier is usually referred to as the *intrinsic*^20,28^ barrier for stalk formation (around ξ_C_ = 0.75). The barrier is similar for S1 and S2, being only marginally lower in S1 (∼15 kJ/mol against ∼17 kJ/mol), which is coherent with the observation of a similar timescale for the stalk formation in unbiased MD simulations. However, the pathway to stalk formation is thermodynamically favored in the **S2** system, for which *state S* is much more stable than in **S1**, resulting in the most thermodynamically stable state. Therefore, our simulations suggest that the stabilization of the NP-induced stalk state due to cholesterol should favor the fusion process, while the stalk instability in the absence of cholesterol should reduce the frequency of NP-driven fusion events.

## DISCUSSION AND CONCLUSIONS

Amphiphilic AuNPs share with fusogenic proteins and peptides the ability to anchor to the hydrophobic core of lipid membranes via a linear hydrophobic ligand. Like SNARE complexes, the same NP can interact with two bilayers simultaneously. NPs and fusogenic protein complexes, though, differ in many respects. MUS:OT NPs retain a quite spherical shape during stalk formation, at variance with the elongated SNARE zippers. MUS:OT NPs penetrate the membrane core with at least part of their bulky gold core, and the stalk forms on top of the NP itself, while SNARE zippers favor stalk formation remaining in the inter-membrane water compartment. Above all, the interactions of MUS:OT nanoparticles with lipid membranes are less specific than those of their biological counterparts. At variance with the terminal segments of SNARE proteins and with fusogenic peptides, the NP ligands neither fold nor undergo significant structural rearrangements while anchoring to the membrane core. The NPs interact non-specifically with divalent ions, contrary to proteins such as synaptotagmin or Munc13, whose fusogenic action is specifically Ca^2+-^triggered.

Such similarities and differences pose an untrivial question – how does the overall physical path to fusion change in the presence of fusogenic nanoparticles? The data presented in this paper allow us to offer some molecular-scale answers to this question, which remains open and challenging in many respects.

The overall barrier for the formation of the first fusion intermediate, the fusion stalk, is composed of two distinct barriers. The first is related to the dehydration barrier, which needs to be overcome to bring the two membranes close to each other. The second barrier is associated with pulling a molecular-scale trigger, such as the protrusion of a lipid tail^20,28^, establishing the first hydrophobic contact between the two juxtaposing membranes.

Previous computational analyses have shown that many different effects can reduce dehydration barriers – ions in solution, presence of charged lipids in the membrane, and effective membrane surface hydrophobicity^28^. Lipids with a spontaneous negative curvature are less efficient at shielding the hydrophobic tails from solvent. Therefore, a smaller head group, such as that of cholesterol, causes a more prominent exposure of lipid tails to the solvent, reducing membrane-membrane repulsion^20,75^. In the presence of MUS:OT nanoparticles, the relevant distance is no longer the distance between the two membranes but rather the distance between the water-exposed ligands of the nanoparticle and the juxtaposed membrane. There is strong evidence, both experimental and computational^76,77^, that MUS:OT nanoparticles are effective at creating, spontaneously, close membrane-membrane contacts. These interactions can be caused by single nanoparticles^36,77^ as well as by large nanoparticle aggregates. Molecular Dynamics simulations show that NP ligand-membrane interactions can be direct or ion-mediated, and in any case, they are short-range interactions, implying that the distance between the water-exposed ligands of NPs and the membrane is often shorter than 1 nm. The observation of spontaneous liposome-liposome adhesion in experimental setups and unbiased MD simulations suggests that NPs can lower dehydration barriers to the order of a few k_B_T, outperforming the effects of lipid composition (e.g., cholesterol content^20^), ionic strength, and short-range, ion-mediated lipid-lipid interactions.

Once the dehydration barrier has been overcome, an intrinsic stalk barrier^28^ needs to be climbed. In the absence of NPs, it has been shown that SNARE transmembrane domains^28^ and cholesterol^20^ decrease the intrinsic stalk barrier and stabilize the metastable stalk state. In the presence of a NP, the ligands extending towards the facing membrane offer a significantly large solvent-accessible hydrophobic surface due to the NP curvature and the disordered and flexible ligand nature. Therefore, a lipid protruding from the facing membrane can easily penetrate this hydrophobic surface. Moreover, the surface of the NP can be crawled by lipids that can contribute to establishing a hydrophobic contact between the two membranes. All these processes in our unbiased MD simulations lead to spontaneous stalk formation. Thus, in the presence of NPs, the intrinsic barrier for the stalk formation is significantly lower than in the absence of NPs^20^, and maybe even lower than in the presence of SNARE transmembrane domains^28^ or whole fusogenic protein complexes^25^.

Due to the rough treatment of electrostatics in the coarse-grained model we used and its intrinsically simplified description of ions, it is difficult to use our computational data to evince any specific role of ions in lowering the intrinsic barrier for the stalk formation in the presence of NPs. Atomistically resolved and coarse-grained models, though, agree^77,78^ on the fact that positively charged ions can promote NP-NP aggregation, both in water and in contact with lipid membranes. The possibility that ion-mediated NP-NP interactions further lower stalk formation barriers is something that will deserve future investigations.

The mechanisms by which nanoparticles induce fusion have been partially elucidated. However, it is not clear yet how the fusion process evolves from the stable, cholesterol-stabilized fusion stalk shown in Figure 6 to pore opening. Pore opening may result from the collective action of two or more nanoparticles, not dissimilarly from what happens with SNARE pins^79^. Ions may also take part in the process in another way by favoring NP-NP aggregation on the surface of the interacting membranes.

Despite their intrinsic aspecificity, we have shown in this paper that the fusogenic activity of small (2-3 nm) amphiphilic AuNPs can be finely tuned by exploiting divalent ions and membrane cholesterol. While the NP-mediated juxtaposition of two bilayers, at distances of the order of a couple of nm, is spontaneous and largely effective even in the absence of ions and cholesterol^77^, these two ingredients concur at lowering the intrinsic stalk barrier, stabilizing the metastable stalk state and possibly, by NP-promoted mechanisms that have not been understood yet, favoring the opening of the fusion pore. These NPs are thus promising for applications requiring *in vitro* controlled membrane fusion, such as loading artificial liposomes or exosomes for drug delivery applications^11^.

