## Supplementary Information for "Cholesterol-containing liposomes decorated with Au nanoparticles as minimal tunable fusion machinery"

Ester Canepa[ac], Davide Bochicchio[a], Paulo Henrique Jacob Silva[b], Francesco Stellacci[b], Silvia Dante[c]*, Giulia Rossi[a]*, Annalisa Relini[a]

a Department of Physics, University of Genoa, 16146, Genoa, Italy
b Institute of Materials Science & Engineering, EPFL, 1015, Lausanne, Switzerland
c Materials Characterization Facility, Istituto Italiano di Tecnologia, 16163, Genoa, Italy

### Materials

1,2-dioleoyl-*sn*-glycero-3-phosphocholine (DOPC, 18:1(Δ9-Cis) PC, >99%), 1,2-dioleoyl-*sn*-glycero-3-phosphoethanolamine-N-(7-nitro-2-1,3-benzoxadiazol-4-yl) (ammonium salt) (18:1 NBD-PE, >99%) and 1,2-dioleoyl-*sn*-glycero-3-phosphoethanolamine-N-(lissamine rhodamine B sulfonyl) (ammonium salt) (18:1 Liss Rhod-PE, >99%) were purchased as lyophilised powders from Avanti Polar Lipids. Cholesterol (chol, ≥99 %, lyophilized powder), chloroform ($CHCl_3$, ≥99.5 %), Tris(hydroxymethyl)aminomethane (Trizma® base, >99.9 %), sodium chloride (NaCl, ≥99.5 %), Sulphorodamine B (acid form) (SRB, dye content 95 %), calcium chloride dihydrate ($CaCl_2 \cdot 2H_2O$, ≥99.0%) and magnesium chloride hexahydrate ($MgCl_2 \cdot 6H_2O$, 99.0–102.0%) were purchased from Sigma Aldrich (Merck). See Canepa et al[1] for all materials used in NP synthesis. All chemicals were used without further purification. Before use, water was always purified with a Milli-Q ultrapure water system (18.2 MΩ × cm resistivity at 25° C; Merck Millipore).

### NP synthesis and experimental characterization

*NP synthesis.* For this study, we used the same monodisperse 2:1 MUS:OT AuNPs employed in our previous work[1]. NP powders were suspended in water (6 mg/mL) and diluted in 2.5 mM Trizma® base and 50 mM NaCl (pH 7.4) before experiments. Both in water and in the experimental buffer, 2:1 MUS:OT AuNPs showed remarkable long-term colloidal stability.

*NP characterization*. Monodisperse 2:1 MUS:OT AuNPs were characterized by Transmission Electron Microscopy (TEM), Nuclear Magnetic Resonance (NMR), Dynamic Light Scattering (DLS), ζ-potential analysis and UV-Vis spectroscopy. Results are shown in Table S1 and Figures S1. Details on TEM and NMR analyses (for core size and ligand ratio characterization, respectively) are described in Canepa et al[1]. For both DLS and ζ-potential measurements, a Malvern Zetasizer Nano ZS instrument (scattered light collected in backscattering at 173° ) was used, whereas UV-Vis measurements were performed using a Jasco V-530 spectrophotometer. DLS analysis was carried out at 25° C in water and in buffer by diluting the NP stock solution to 0.3 mg/mL[1] and 0.03 mg/mL, respectively. The NP ζ-potential was also analyzed at 25° C in water and in buffer, but with samples diluted to 0.07 mg/mL[1]. Finally, UV-vis characterization was performed in buffer at 0.03 mg/mL. In all cases, NP dispersions were gently sonicated in an ultrasonic bath for a few minutes before analysis.

**Table S1.** Characterization of monodisperse 2:1 MUS:OT AuNPs used in the experiments.

| TEM | $^1$H NMR | DLS – hydrodynamic diameter (nm) | | ζ-potential (mV) | |
|---|---|---|---|---|---|
| core diameter (nm) | OT mol % | water | buffer [c] | water | buffer [c] |
| 2.4 ± 0.4 [a] | 32 ± 5 [b] | 5.6 ± 1.7 [d] | 9.4 ± 0.4 [d] | –46 ± 5 [d] | –34 ± 3 [d] |

a. mean ± one standard deviation.

b. mean ± error calculated using Student's statistics (95 % confidence level, N=3: three NMR spectra repeated on separately weighed NP samples).

c. 2.5 mM Trizma® base and 50 mM NaCl (pH 7.4; 25° C).

d. mean ± error calculated using Student's statistics (95 % confidence level, N=12÷18); hydrodynamic diameters were averaged from number distributions (N %).

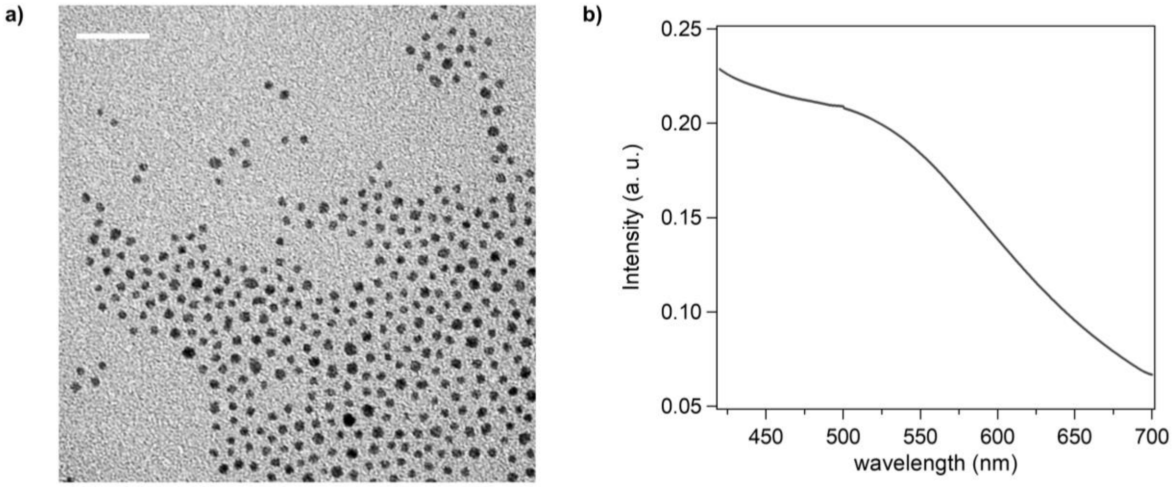


**Figure S1. a)** Bright-field TEM image of monodisperse 2:1 MUS:OT AuNPs used in this study (scale bar 20 nm). Core diameter: 2.4 ± 0.4 nm (mean ± one std dev). **b)** UV-Vis spectrum of the 2:1 MUS:OT AuNPs shown in (a) recorded in the experimental buffer (0.03 mg/mL) using a Suprasil® quartz cuvette (Hellma®, semi micro, light path 10 × 4 mm). This analysis shows the characteristic Localized Surface Plasmon Resonance (LSPR) peak of these NPs at ~520 nm[2].

The DLS (hydrodynamic size) and ζ-potential results reported in Table S1 indicate that 2:1 MUS:OT AuNPs are colloidally stable under the experimental conditions used in our investigations (pH 7.4, 25° C). As shown in Figures S2, these NPs undergo marked aggregation following the addition of 2 mM $Ca^{2+}$ during membrane fusion experiments (pH 7.4, 25° C).

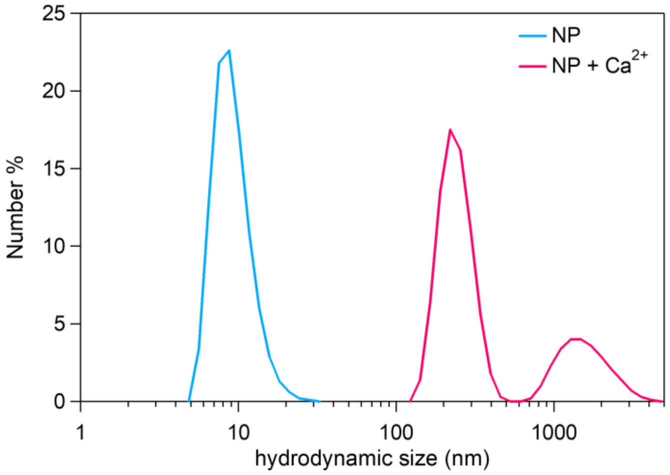


**Figure S2.** Hydrodynamic diameter (measured by DLS) of 2:1 MUS:OT AuNPs in the experimental buffer (2.5 mM Trizma® base and 50 mM NaCl; pH 7.4, 25° C) before and after $Ca^{2+}$ addition (2 mM). The blue peak in the absence of calcium ions corresponds to 9.4 ± 0.4 nm (Table S1).

### NP structure and computational model

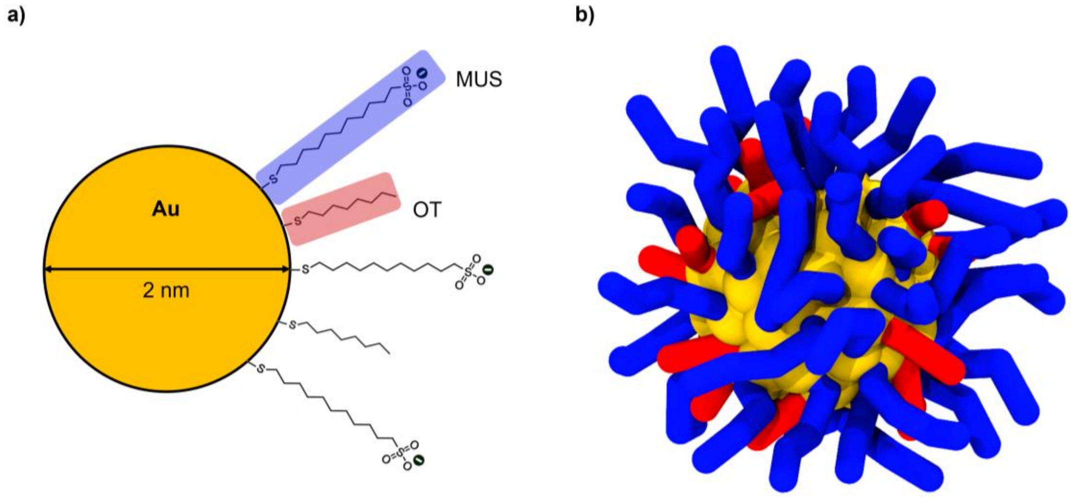


**Figure S3.** Structure and coarse-grained model of the amphiphilic MUS:OT NPs used in this study. **a)** Chemical structure of the two alkylthiol ligands (MUS and OT) that functionalise the AuNP surface. **b)** Coarse-grained representation of the same MUS:OT NP, in which the gold core is in yellow, the MUS ligands in blue, and the OT ligands in red.

### Lipid vesicle preparation and experimental characterization

*Lipid vesicle preparation*. DOPC vesicles with 0, 10, 20, 30 and 40 mol % chol were prepared according to the procedure described in our previous work[1]. Specifically, vesicles were extruded 25 times in 2.5 mM Trizma® base and 50 mM NaCl (pH 7.4)[3] using the Avanti Mini-Extruder (Avanti Polar Lipids) with a 0.1 µm pore diameter polycarbonate membrane (Nuclepore filters, Whatman). For SAXS analysis only, a membrane with a pore diameter of 0.05 µm was employed. For QCM-D and fluorescence measurements, lipid vesicles were prepared at 2 mg/mL, whereas a 5-fold higher concentration was used for SAXS experiments. For lipid mixing assays, we labeled the vesicle bilayers by adding 0.5 mol % 18:1 NBD-PE and 0.5 mol % 18:1 Rhod-PE[4]. Stock solutions (1 mg/mL) of the fluorescent lipids were previously prepared in $CHCl_3$. For content mixing assays, vesicles were extruded using a self-quenched SRB solution (50 mM)[3,5] prepared in water and containing 2.5 mM Trizma® base (pH 7.4). At this concentration, SRB is known to undergo >95% self-quenching[6]. SRB-loaded vesicles were filtrated from the non-encapsulated dye by the minicolumn centrifugation technique[7] as described in Canepa et al[8]. During filtration, the external SRB solution was replaced by the experimental buffer containing 2.5 mM Trizma® base and 50 mM NaCl (pH 7.4). All vesicle suspensions were stored at 4° C and used within a few days; batches containing fluorescent probes were also protected from light till further use.

*Lipid vesicle characterization*. To characterize the hydrodynamic size of lipid vesicles, DLS measurements were performed at room temperature using the same instrument reported for NP characterization. Measurements were carried out right after extrusion, without vesicle dilution. In the case of SRB-containing vesicles, DLS analysis was performed after vesicle filtration. Vesicle size results are reported in Figure S4.

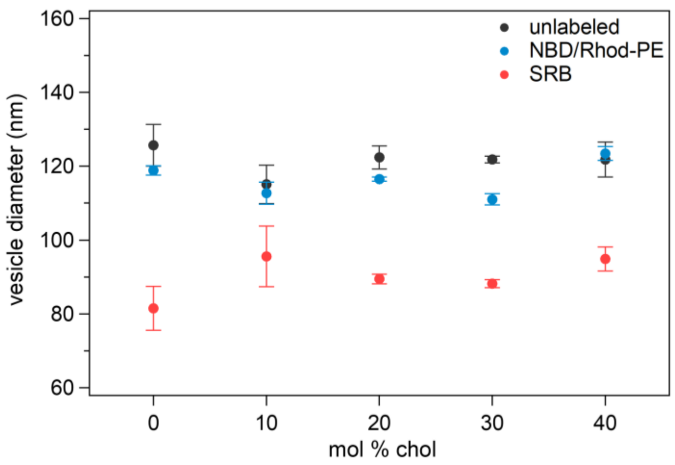


**Figure S4.** Hydrodynamic diameters (mean ± one std dev) of lipid vesicles at varying membrane cholesterol concentration (unlabeled vesicles, vesicles labeled with NBD-PE and Rhod-PE, and vesicles containing self-quenched SRB). Data obtained from intensity distributions (N=12).

### QCM-D experiments

The dissipative QCM investigation was carried out using the same setup and protocol described in Canepa et al[1]. Viscoelastic supported vesicle layers (SVLs) were deposited onto the surface of a gold-coated QCM sensor and used to quantify the passive uptake of amphiphilic NPs in fluid lipid membranes with increasing cholesterol content. Further QCM-D investigations were performed to study vesicle fusion promoted by membrane-embedded NPs in the presence of $Ca^{2+}$ and $Mg^{2+}$ ions. All experiments were performed at 25° C in 2.5 mM Trizma® base and 50 mM NaCl (pH 7.4) in a QCM-Z500 microbalance (KSV Finland LLC) equipped with a thermostated flow chamber.

*Sample preparation*. Freshly extruded vesicles were diluted in the experimental buffer (0.25 mg/mL) and let to equilibrate at 25° C in the QCM pre-chamber for 10 min. Vesicles were then injected into the QCM chamber to initiate the formation of SVLs (Figure 1a, main text). Before NP addition, each SVL was gently rinsed with fresh buffer thermostated at 25° C to remove excess vesicles (Figure 1a, main text). Amphiphilic NPs were then diluted in buffer (0.01 mg/ml), gently sonicated for a few minutes in an ultrasonic bath and finally inserted into the QCM pre-chamber. After 10 min of equilibration at 25° C, NPs were injected into the main chamber and allowed to interact with the preformed SVL for at least 1 h (Figure 1a, main text). The SVL-NP complex was finally rinsed with buffer before the end of the recording. In general, no mass loss was detected after rinsing indicating a stable NP insertion within the vesicle layer (Figures 1a and 5a, main text).

For membrane fusion experiments (Figure 5a,b, main text), we waited only ~900 s of SVL-NP incubation before removing NPs that had not passively penetrated the vesicle bilayer. Fresh vesicles with the same cholesterol content as the SVL were then diluted in buffer (0.05 mg/mL), mixed with $Ca^{2+}$ or $Mg^{2+}$ ions and placed in the QCM chamber to interact with the rinsed SVL-NP complex. Both $Ca^{2+}$ or $Mg^{2+}$ were diluted in buffer to 2 mM using aqueous stock solutions of $CaCl_2$ or $MgCl_2$ (0.17 M). After insertion of vesicles and divalent ions, the sample was monitored for approximately 40÷60 min before the final rinse with buffer. In all cases, the final rinsing of post-fusion samples did not result in the loss of material from the sensor surface (Figure 5b, main text). Control membrane fusion experiments were conducted without adding NPs to the SVL or by adding, after NP uptake, only the divalent ions (i.e. without vesicles, Figure S14).

*Data processing*. As in our previous work[1], QCM-D data were processed in the assumption of a modified Sauerbrey model to include the change in viscosity typical of viscoelastic layers:

$$\Delta \mathrm{m} = -C_f \frac{\Delta \mathrm{f}}{n^{\frac{1}{2}}} \tag{1}$$

where $n$ is the overtone number at which the QCM-D quartz crystal sensor is driven, $\frac{\Delta \mathrm{f}}{n^{\frac{1}{2}}}$ (or $\Delta f_{norm}$) is the observed frequency shift ($\Delta$f, Hz) normalized by $n^{\frac{1}{2}}$ instead of the canonical $n$ used for rigid films, $C_f$ is the sensitivity coefficient of the quartz crystal sensor (17.7 ng/($cm^2$ • Hz)), and $\Delta \mathrm{m}$ is the change in mass per unit area (ng/$cm^2$) of the piezoelectrically active surface (78.5 $mm^2$).

This data processing allows to convert the frequency shifts ($\Delta$f) detected during the recording (Figures 1a and 5a, main text) into the mass changes on the sensor surface corresponding to the formation of viscoelastic SVLs (i.e. $m_{SVL}$) and the following uptake of NPs (i.e. $m_{NP}$). These values were subsequently used to monitor over time the percentage change in SVL mass during passive NP incorporation:

$$\Delta \mathrm{m}_{SVL}(t)\ (\%) = \frac{m_{NP}(t)}{m_{SVL}} \cdot 100 \tag{2}$$

where $m_{SVL}$ was calculated after rinsing the preformed SVL before the addition of NPs (Figures 1b, main text).

This data processing has already proven to be suitable and reliable for identifying mass percentage changes in viscoelastic films, such as SVLs[1]. The same data processing was also performed to report the $\Delta \mathrm{m}_{SVL}(t)$ (%) traces due to the membrane fusion processes shown in Figure 5b (main text). Here, mass changes recorded after the rinsing of excess NPs and the addition of vesicles/$Ca^{2+}$ (or vesicles/$Mg^{2+}$) were considered instead of those due to NP adsorption. All percentage mass changes reported in the main text (Figure 1b and Figure 5b) were plotted by averaging the results obtained from the different overtones (n=3, 5, 7, 9, 11).

### Fluorescence-based membrane fusion assays

All fluorescence measurements were performed in a Suprasil® quartz cuvette (Hellma®, semi micro, light path 10×4 mm) using a Fluorolog-3 spectrofluorometer (Horiba Jobin-Ivon) equipped with temperature control of the cell holder. Experiments were conducted under continuous stirring at 25° C in 2.5 mM Trizma® base and 50 mM NaCl (pH 7.4). Aliquots from an aqueous stock solution of $CaCl_2$ or $MgCl_2$ (0.17 M) were used to trigger membrane fusion.

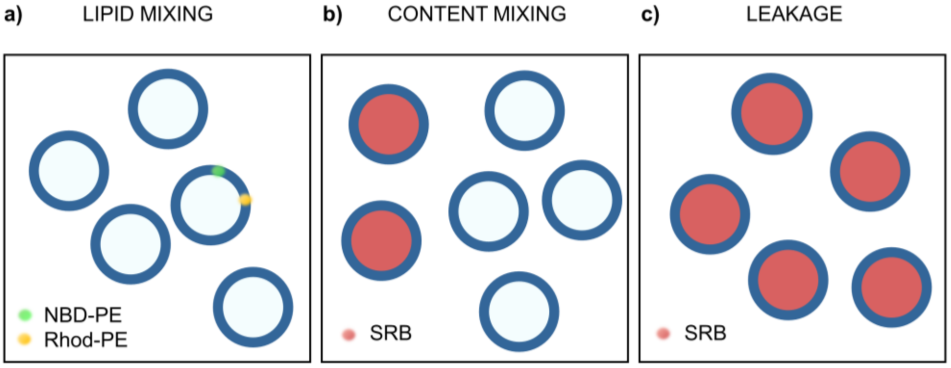


**Figure S5.** Schematic drawing illustrating the vesicle systems used to test membrane fusion by fluorescence assays.

*Lipid mixing*. Non-fluorescent vesicles and vesicles labeled with NBD-PE and Rhod-PE were mixed in a 4:1 molar ratio to achieve a lipid concentration of 0.32 mM in 850 μL total volume (Figure S5a). Samples were excited at $\lambda_{exc}$=470 nm, the characteristic excitation peak of the donor fluorophore (NBD-PE). The fluorescence of NBD-PE was registered over time at $\lambda_{em1}$=530 nm, while that of the acceptor fluorophore Rhod-PE ($\lambda_{exc}$=540 nm) at $\lambda_{em2}$=585 nm. Regardless of the percentage of membrane cholesterol, vesicles and NPs were initially allowed to incubate in buffer at a molar ratio $(\frac{mol_{lipids}}{mol_{NPs}})$ of 580 to promote passive NP uptake into the membrane. This ratio corresponds to a NP concentration in cuvette equal to 0.03 mg/mL, the same as used for DLS and UV-Vis characterization in buffer. After 5 min of NP-vesicle incubation, 10 μL of calcium stock solution (see above) was added to reach a cuvette concentration of 2 mM[3]. The fluorescence emission of lipid-bound FRET dyes NBD and rhodamine was subsequently monitored for 30 min before the end of the recording. The same protocol was also adopted for experiments in which lower $Ca^{2+}$ concentrations were tested (100 μM÷2 mM; Figure 3a-c, main text). Figure S7 shows the emission spectra of a representative lipid mixing experiment conducted for this study (2 mM $Ca^{2+}$).

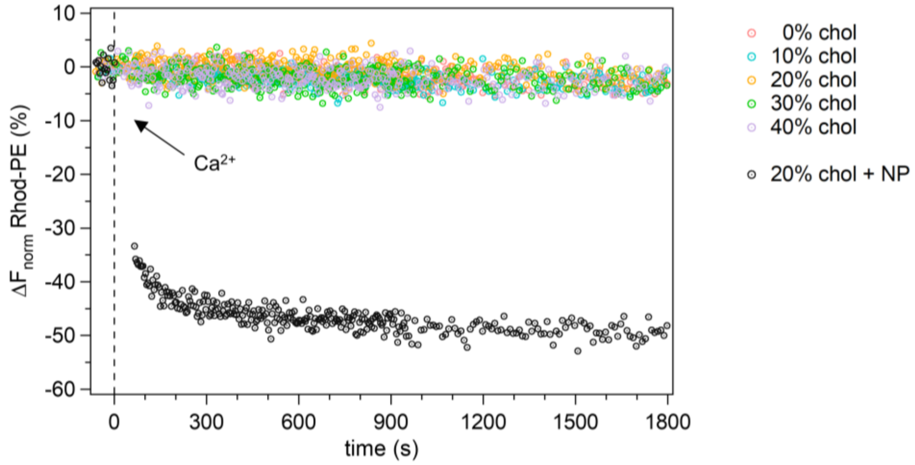


**Figure S6.** Control lipid mixing assays without NPs. Normalized reduction (%) of Rhod-PE signal before and after calcium addition to vesicles incubated or not with NPs (4 s sampling interval before 900 s, 10 s after). In the case of NP-vesicle incubation (5 min), only one representative experiment is shown (20 mol % chol). In general, $Ca^{2+}$ addition induced no change in vesicle fluorescence emission in the absence of NPs.

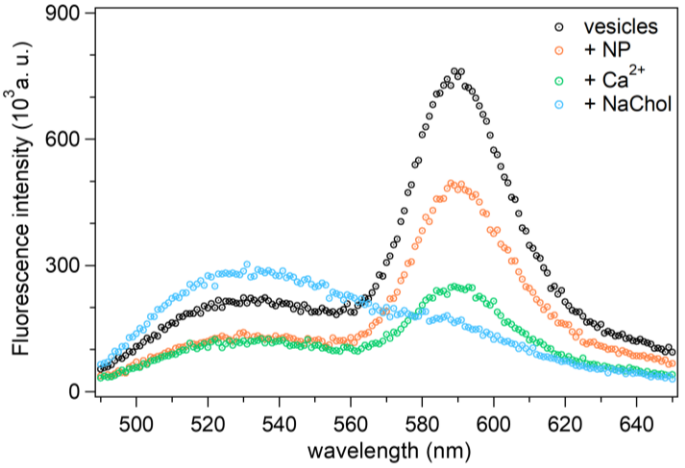


**Figure S7.** Emission spectra corresponding to typical steps of a lipid mixing assay involving unlabeled vesicles and vesicles labeled with lipid-bound FRET dyes (20 mol % chol). NBD-PE (donor) has a maximum emission at 530 nm, while Rhod-PE (acceptor) at 585 nm. Step 1: vesicles alone (black curve). Step 2: vesicles incubated with NPs for 5 min (orange curve). As shown by the orange curve, 2÷3 nm 2:1 MUS:OT AuNPs ($\lambda_{LSPR}$~520 nm, Figure S1b) significantly quench the emission of the donor probe, and consequently also that of the acceptor. Step 3: NP-vesicle complex after $Ca^{2+}$ addition (2 mM, green curve). Despite the strong quenching of the donor emission by NPs, the reduction in Rhod-PE emission observed in the green curve indicates that a spatial separation between the acceptor and donor probe has occurred due to lipid mixing. This representative experiment also includes an additional spectrum (blue curve) recorded after vesicle rupture by detergent addition (20 µL sodium cholate, NaChol, 0.25 mg/mL stock solution in water). Vesicle solubilization is expected to maximize lipid mixing, significantly reducing the proximity-dependent FRET signal. The disappearance of the Rhod-PE peak and the increase of the NBD-PE emission therefore confirms the correct functioning of FRET in our lipid system.

To verify whether the FRET signal recorded during lipid mixing experiments derived from the inner or outer leaflet of the membrane bilayer, we tested the vesicle system shown in Figure S5a with a quencher of NBD fluorescence. Specifically, we added small volumes of a fresh aqueous solution of sodium dithionite (≥ 85.0%, Sigma Aldrich-Merck) (0.86 M) to reach a final concentration of 70 mM (Figure S8). Since sodium dithionite penetrates very slowly into intact phospholipid bilayers, within minutes of its addition it is only able to quench the fluorescence of NBD-labeled lipids that are exposed to the external medium[9,10]. The dithionite quenching results shown in Figure S8 report no change in the FRET signal after sequential quencher additions, thus revealing that the NBD-PE probe is mostly located on the inner surface of our vesicles. This evidence provides a simple method to demonstrate that the lipid mixing assessed in this study by FRET measurements (Figures 2b and 3d, main text) occurs predominantly in the inner vesicle leaflet.

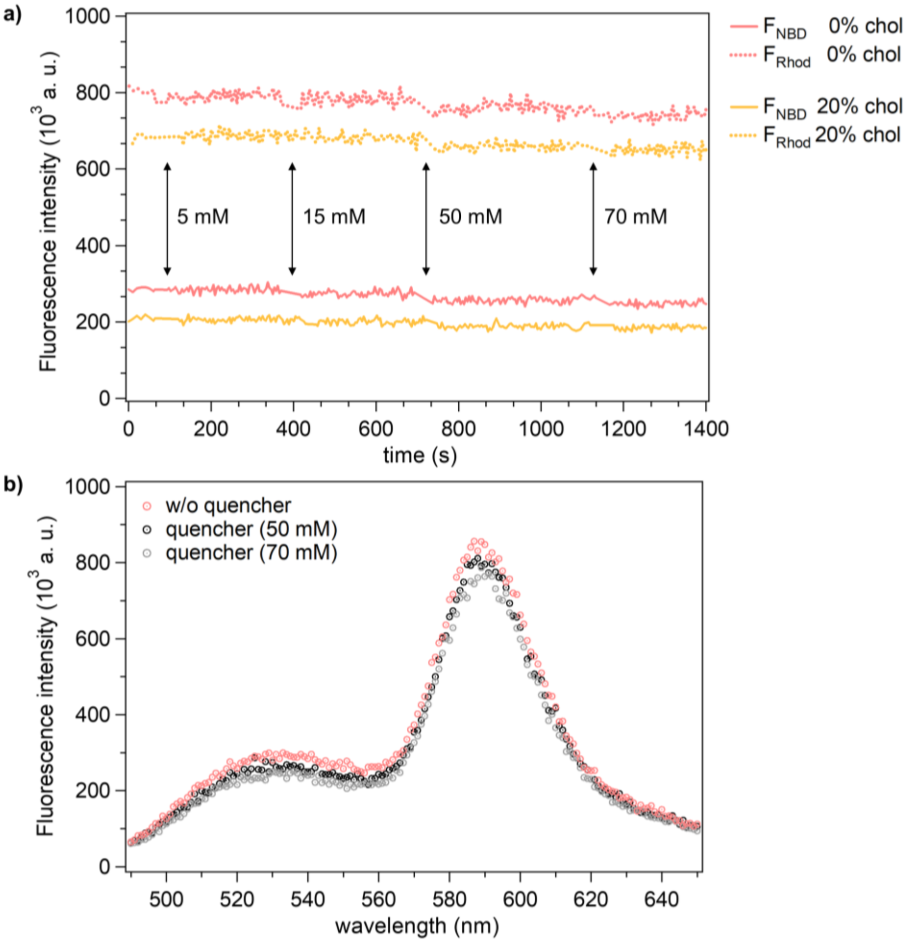


**Figure S8. a)** Monitoring of FRET signal during four sequential additions of sodium dithionite to lipid vesicles (0.32 mM lipid concentration, 850 μL total volume) (4 s sampling interval). Representative traces of NBD-PE and Rhod-PE emission are reported in the presence (20 mol %) and absence of membrane cholesterol. In general, only at high quencher concentrations in the external buffer (i.e. 50 mM) a slight decrease in the FRET signal was recorded. Further dithionite additions did not lead to further FRET quenching. **b)** Emission spectra corresponding to the 0 mol % cholesterol traces shown in (a). The black curve at 50 mM dithionite and the grey curve at 70 mM dithionite were recorded ~ 2 min and ~ 4 min after addition of the quencher, respectively.

*Content mixing*. Non-fluorescent vesicles and vesicles loaded with self-quenched SRB were mixed in a 2:1 molar ratio to achieve a total lipid concentration of 0.32 mM in 850 µL total volume (Figure S5b). During experiments, the fluorescence emission of SRB, excited at $\lambda_{exc}$=565 nm, was continuously recorded at $\lambda_{em}$=586 nm. As for lipid-mixing assays, at all cholesterol percentages vesicles and NPs were let to interact 5 min in buffer at a molar ratio $(\frac{mol_{lipids}}{mol_{NPs}})$ of 580 before calcium addition (2 mM final concentration)[3]. Sample fluorescence was monitored for 30 min before the addition of 20 µL of an aqueous stock solution of sodium cholate (≥99%, Sigma Aldrich-Merck) (0.25 mg/mL) to induce complete leakage of the inner dye (Figure S10). The same protocol was also adopted for experiments in which the effect of $Mg^{2+}$ ions, instead of $Ca^{2+}$, was tested (Figure 5c, main text).

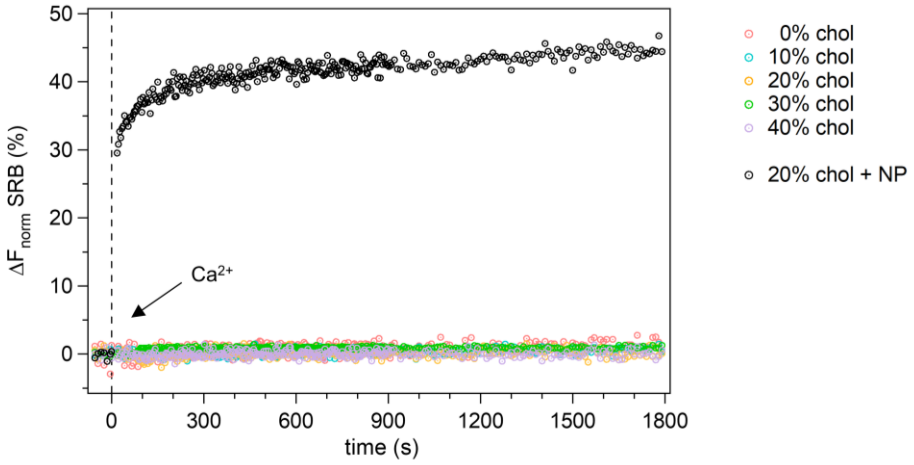


**Figure S9.** Control fluorescence assays without NPs. **a)** Normalized reduction (%) of Rhod-PE signal and **b)** normalized increase (%) in SRB signal before and after calcium addition to vesicles incubated or not with NPs (4 s sampling interval before 900 s, 10 s after). In the case of NP-vesicle incubation (5 min), only one representative experiment is shown (20 mol % chol). In general, $Ca^{2+}$ addition induced no change in vesicle fluorescence emission in the absence of NPs.

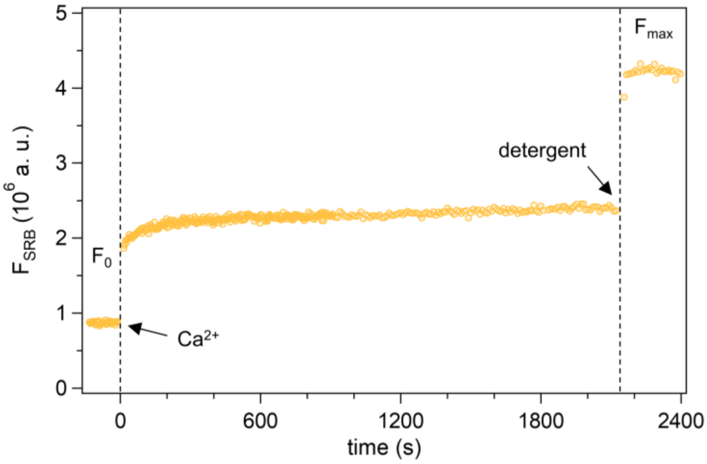


**Figure S10.** Typical content mixing trace before data normalization: representative case at 20 mol % chol with 5 min NP-vesicle incubation (4 s sampling interval before 900 s, 10 s after). $F_{0, SRB}$ is the mean fluorescence level of the self-quenched SRB immediately before calcium addition, while $F_{max, SRB}$ is the mean fluorescence level of the dye after the detergent-induced vesicle rupture. These values are used to normalize the time-dependent SRB fluorescence emission monitored during the fusion event (see Equation 4).

*Leakage.* Leakage experiments were performed using the same protocol as described for content mixing assays, but with only SRB-loaded vesicles (0.32 mM in 850 µL total volume, Figure S5c). In this lipid system, increases in the SRB emission, such as that shown in Figure S11, can only be attributed to dilution of the probe with the external medium due to vesicle content leakage during the fusion event.

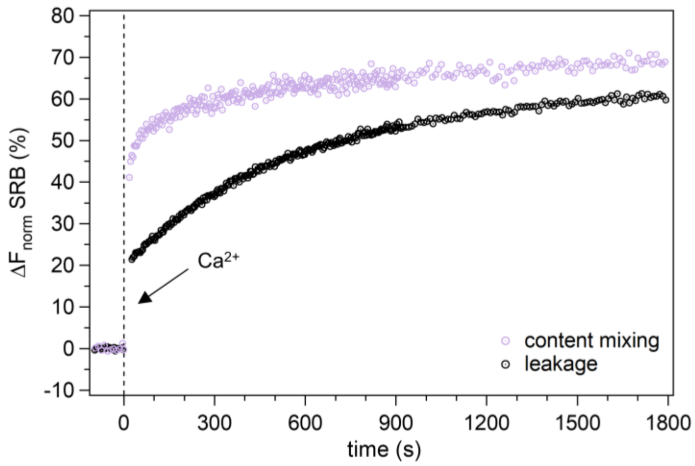


**Figure S11.** Representative leakage experiment (40 mol % chol) compared with the content mixing trace at the same membrane composition (4 s sampling interval before 900 s, 10 s after). In both assays, $Ca^{2+}$ was added after 5 min of NP-vesicle incubation ($\frac{mol_{lipids}}{mol_{NPs}}$=580). This result shows that NP-mediated membrane fusion, and the corresponding rapid content mixing, occurs simultaneously with a slower, partial leakage into the external medium of the vesicle contents involved in the fusion event.

*Data processing*. For FRET-based lipid mixing assays, the reduction in the fluorescence emission of the acceptor lipid probe (Rhod-PE) due to the increase in the average spatial separation from the donor lipid probe (NBD-PE) was calculated as:

$$\Delta \mathrm{F}_{norm} Rhod\ (\%) = \frac{F_{Rhod}(t) - F_{0,Rhod}}{F_{0,Rhod}} \cdot 100 \tag{3}$$

where $\Delta \mathrm{F}_{norm} Rhod\ (\%)$ is the normalized percentage decrease in Rhod-PE fluorescence, $F_{Rhod}(t)$ is the Rhod-PE time-dependent fluorescence and $F_{0,Rhod}$ the mean fluorescence level immediately before calcium addition. A similar equation can be used to normalize the percentage increase in NBD emission. Since 2:1 MUS:OT AuNPs ( $\lambda_{LSPR}$~520 nm, Figure S1b) considerably quench the NBD signal at 530 nm[3] – making changes in its signal less appreciable (Figure S7) – in this paper we discuss the lipid mixing experiments focusing on the most noticeable decrease in Rhod-PE emission.

For content-mixing and leakage assays based on SRB dilution, fluorescence data were normalized as follows:

$$\Delta \mathrm{F}_{norm} SRB\ (\%) = \frac{F_{SRB}(t) - F_{0,SRB}}{F_{max,SRB} - F_{0,SRB}} \cdot 100 \tag{4}$$

where $\Delta \mathrm{F}_{norm} Rhod\ (\%)$ is the normalised percentage increase in SRB emission, $F_{SRB}(t)$ is the time-dependent SRB fluorescence, $F_{0,SRB}$ the mean SRB fluorescence level immediately before calcium addition and $F_{max,SRB}$ the maximum SRB fluorescence after detergent injection (sodium cholate) (Figure S10).

### SAXS experiments

SAXS experiments were carried out using a Malvern PANalytical third generation Empyrean multipurpose platform. Cu ka radiation (l=1.54 Å) was used (40 kV, 45 mA). 1D-SAXS measurements were carried out in a vacuum path chamber (Scatter $X^{78}$) using a beam with line collimation and a $GaliPIX^{3D}$ detector. Scans were performed in the 2q region region 0.06° –5.00° with stepsize 0.014° . Quartz capillary (Hilgenberg, DE) with a 1mm diameter (100 mL volume) were used.

*Sample preparation*. Background measurement of the experimental buffer (2.5 mM Trizma® base and 50 mM NaCl; pH 7.4) was performed in each of the capillary used. Vesicle samples were prepared as described above. Vesicles were extruded using 50 nm polycarbonate filters to meet the requirement of instrumental resolution at low angle. The vesicle concentration was 10 mg/mL. DOPC samples containing 0, 20, 30, and 40 mol % cholesterol were prepared. Diluted NP were added to the vesicles to avoid scattering from aggregated NPs upon $Ca^{2+}$ ions addition, as known by DLS measurements (Figure S2). To this aim, a molar ratio ($\frac{mol_{lipids}}{mol_{NPs}}$) of 16240 was adopted, corresponding to ~1 NP/vesicle and 28 times lower than that used for fluorescence assays. $Ca^{2+}$ was added at 2 mM concentration (from $CaCl_2$ 0.17 M), as in all previous experiments.

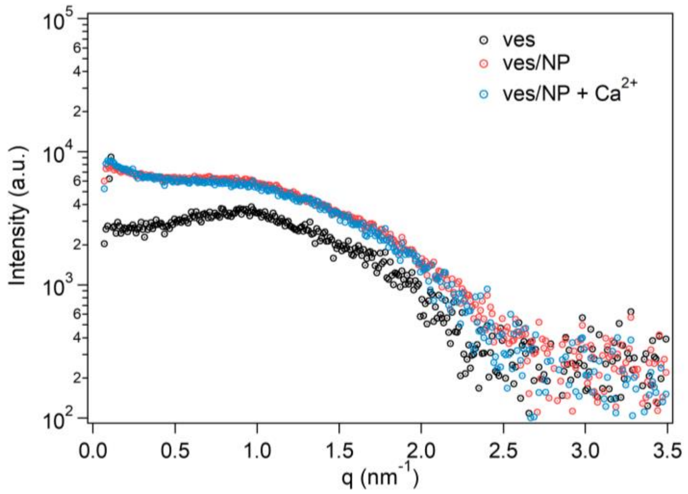


**Figure S12.** SAXS curves of pure DOPC vesicles (0 mol % chol) before (black circles) and after (red circles) the incubation with NPs. The subsequent administration of $Ca^{2+}$ ions (2 mM) does not change the scattering profile in the observed q range (blue circles). Before interaction with NPs the vesicle radius is 25 nm. The behavior is similar to that reported in the main text for 20 mol % chol (Figure 4a).

*Data fitting.* The data analysis was performed using the EasySAXS software (Malvern-PANalytical). After background subtraction, the experimental curves were fitted using a spherical shape with radius $r$ and a concentric core-shell structure consisting of three shells, i.e. the inner and outer lipid headgroup regions and the hydrophobic bilayer core (Figure S13a). Each shell was therefore described by 3 parameters: thickness $t$, electron density contrast $Dr$, and polydispersity. The core contrast was set to zero (the core material is the same as the medium surrounding the particles). The contrast of each shell is defined as the contrast of the shell with respect to the core. The parameters obtained from the best fits are summarized in Table S2. The best fits of the experimental curves are reported in the insets of Figure 4a-c of the main text (20÷40 mol % chol) and in Figure S12 (0 mol % chol). For pure vesicle samples, at all cholesterol molar percentage the fit procedure with spherical particles returned $r$ values in perfect agreement with the expected dimension (50 nm diameter, see Figure 4d of the main text) and a symmetric structure of the lipid bilayer (Table S2). Adding the NPs, the size of the vesicles increased (Figure 4d) and, at the same time, the structure of the shell was affected. Due to the low S/N in the high-q region, we prefer not to speculate about the rearrangements occurring internally the bilayer.

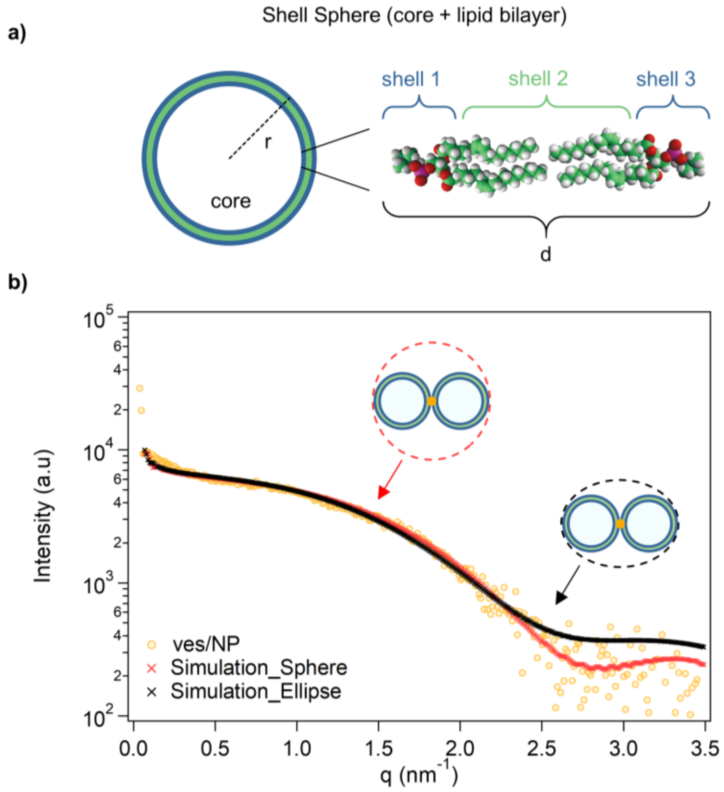


**Figure S13.** Fitting of SAXS experimental curves. **a**) Sketch of the spherical model used for the SAXS fitting procedure. A sphere with radius $r$ and a 3 layers shell, corresponding to the different regions of the lipid bilayer, was used. **b**) In the case of vesicles samples incubated with NPs, an alternative ellipsoidal model was also used.

Scattering curves obtained from vesicles incubated with NPs were also fitted by assuming an ellipsoid with core-shell structure as the shape for the docked vesicles. The best fit obtained in this geometry, with a lipid bilayer with the same structure of the spherical model, is shown in Figure S13b for the representative case of 40 mol % chol. The core of the equatorial semi-axes of the ellipsoid was in this case 60 nm and the ellipsoidal anisotropy ratio n=0.86. The simulated curve was very similar to the one obtained with a spherical symmetry, especially in the low-q region. For sake of simplicity, we reported in the main text the spherical model only.

**Table S2.** Structural parameters obtained for the three layers of the spherical core-shell model ($t$ is the layer thickness and ρ the electron density in arbitrary units).

| | | Chol 0% | | Chol 20% | | Chol 30% | | Chol 40% | |
|---|---|---|---|---|---|---|---|---|---|
| | | ρ (a.u) | $t$ (nm) | ρ (a.u) | $t$ (nm) | ρ (a.u) | $t$ (nm) | ρ (a.u) | $t$ (nm) |
| ***ves*** | *layer 1* | 0,75 | 0,9 | 0,8 | 1,29 | 0,9 | 1,2 | 0,88 | 0,9 |
| | *layer 2* | –0,5 | 2,4 | –0,65 | 2,4 | –0,75 | 2,45 | –0,45 | 2,4 |
| | *layer 3* | 0,75 | 0,9 | 0,8 | 1,24 | 0,9 | 1,2 | 0,4 | 0,9 |
| ***ves/NP*** | *layer 1* | 0,8 | 0,9 | 0,75 | 0,9 | 0,75 | 0,9 | 0,67 | 0,9 |
| | *layer 2* | –0,4 | 2,4 | –0,45 | 2,4 | –0,45 | 2,4 | –0,45 | 2,4 |
| | *layer 3* | 0,55 | 0,9 | 0,45 | 0,5 | 0,45 | 0,9 | 0,45 | 0,5 |
| ***ves/NP/$Ca^{2+}$*** | *layer 1* | 0,89 | 0,9 | 0,75 | 0,9 | 0,75 | 0,9 | 0,75 | 0,9 |
| | *layer 2* | –0,4 | 2,4 | –0,45 | 2,4 | –0,45 | 2,4 | –0,45 | 2,4 |
| | *layer 3* | 0,2 | 0,9 | 0,8 | 0,6 | 0,45 | 0,5 | 0,45 | 0,47 |

### Further control experiments on membrane fusion promoted by NPs (QCM-D and fluorescence)

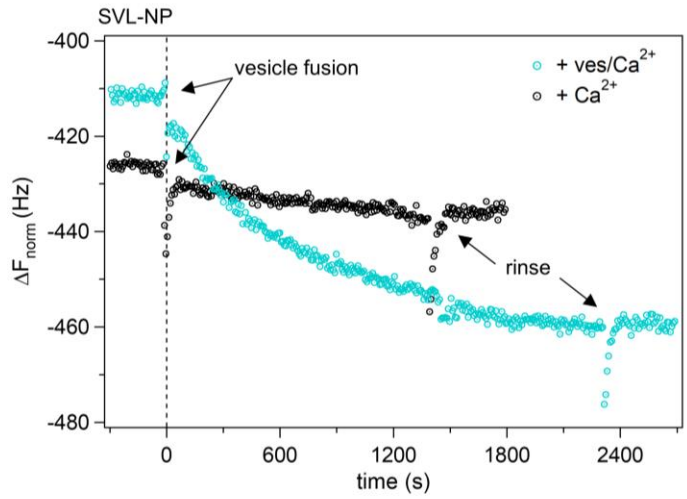


**Figure S14.** Control experiment for studying membrane fusion in SVLs using QCM-D (pH 7.4, 25° C). The traces in the figure refer to an SVL containing 10 mol % chol. The shift in normalized resonance frequency ($\Delta f_{norm}$) recorded after the addition of divalent ions (2 mM) diluted in fresh vesicle buffer (0.05 mg/mL, blue curve) is significantly larger than that observed after the addition of divalent ions alone (2 mM, black curve). The additions (t=0 s) were carried out a few minutes after abundant rinsing of the SVL-NP complex; the sensor resonance frequency was monitored during the entire experiment in real time (7 s sampling interval). The comparison of the two traces shown reveals that, due to the physical constraints imposed by the geometry of the SVL, the study of vesicle fusion on the surface of the QCM sensor is clearly favored when divalent ions mixed with free vesicles are inserted into the QCM chamber.

Therefore, this protocol was preferred to conduct the QCM-based membrane fusion study performed in this work (Figure 5a,b, main text). The plots above refer to the insertion of $Ca^{2+}$ ions, but quite similar results were obtained with $Mg^{2+}$. In all cases, rinsing the SVL-NP complex after fusion did not result in the loss of material from the sensor surface (see also Figure 5b, main text).

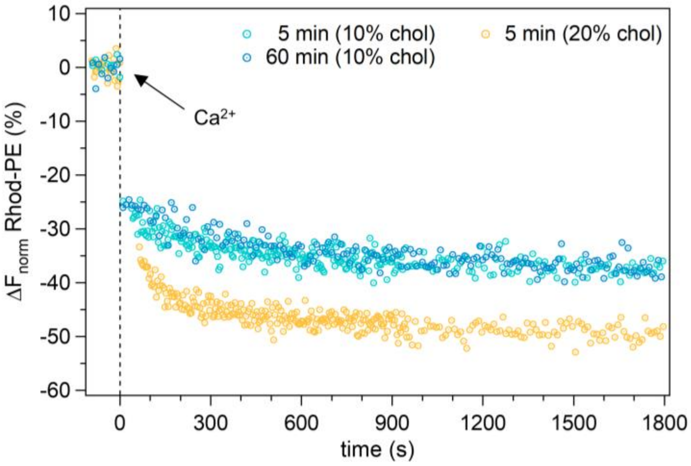


**Figure S15.** Normalized reduction (%) of Rhod-PE signal after addition of $Ca^{2+}$ ions (2 mM). Experiments with vesicles containing 10 mol % chol incubated with NPs for 5 min and 1 h are compared with vesicles containing 20 mol % chol after 5 min incubation. We prolonged the incubation time to 10 mol % chol vesicles as this is the only cholesterol percentage that allows a clear increase in the vesicle uptake of NPs between 5 min and 1 h (Figure 1b, main text). Overall, the synergistic contribution of membrane cholesterol content appears to play a predominant role in promoting membrane fusion compared to the amount of NPs incorporated into the bilayer. Furthermore, the comparison of the curves at 10 mol % chol suggests that the small amount of amphiphilic MUS:OT AuNPs incorporated into the bilayer in the very first minutes of incubation is sufficient to saturate the lipid mixing capacity of the tested vesicles.

### MD simulations

*Definition of* $\xi_C$. The stalk formation is a complicated collective process involving the concerted movement of many molecules and a deep structural rearrangement of the lipid bilayers involved. It is therefore not trivial to define a collective variable (CV) able to describe the relevant minima and barriers along the transition path. Inspired by the work by Hub,[11] exploiting the Plumed plugin,[12] we constructed our CV, $\xi_C$ , defining a cylinder above the NP, with its axis parallel to the membrane normal (z axis) and divided into 8 slices of 0.3 nm of thickness and 2.5 nm of radius (Figure S16).

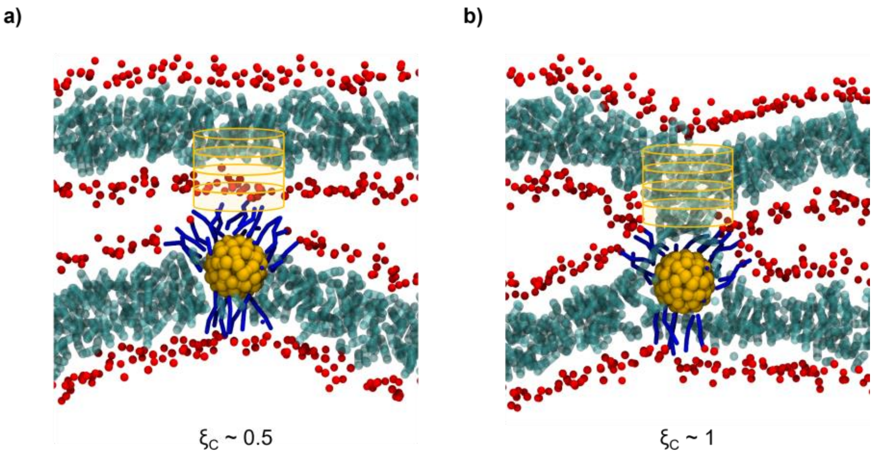


**Figure S16.** Representation of the cylinder used to define $\xi_C$ (only 4 of the 8 slices are shown for clarity). a) The cylinder is placed above the NP core (in yellow), at a distance at which, in the adsorbed state, it is only partially filled with hydrophobic lipid tails (in cyan), and $\xi_C$ is around 0.5. b) When the stalk is formed, the cylinder becomes full of lipid tails and $\xi_C$ approaches 1.

In our unbiased simulations, the stalk always formed above the NP. Therefore, we set the cylinder center in the membrane plane (xy) to coincide with the NP center of mass and added an offset of 1.8 nm along z between the NP COM and the bottom slice of the cylinder. We counted the number $N_i$ of apolar lipid tail beads in each slice $i$ thanks to the Plumed INCYLINDER multicolvar (DIRECTION=Z, RADIUS={TANH R_0=2.5}, SIGMA=0.1), and transformed this value in a filling factor $f_i$ , which varies from 0 to 1, according to the following formula:

$$f_i = 1 - e^{-\frac{N_i}{5}} \tag{5}$$

Then, $\xi_C$ is calculated as the average filling factor over all the slices:

$$\xi_C = \frac{1}{8}\sum_{i=1}^{8} f_i \tag{6}$$

In Figure S17 we report the analysis of two simulation runs (**S1** and **S2** systems) as an example, showing $\xi_C$ as a function of simulation time. It can be noticed how the variable is well suited to identify the stalk formation and the different stalk stability in the **S1** and **S2** systems.

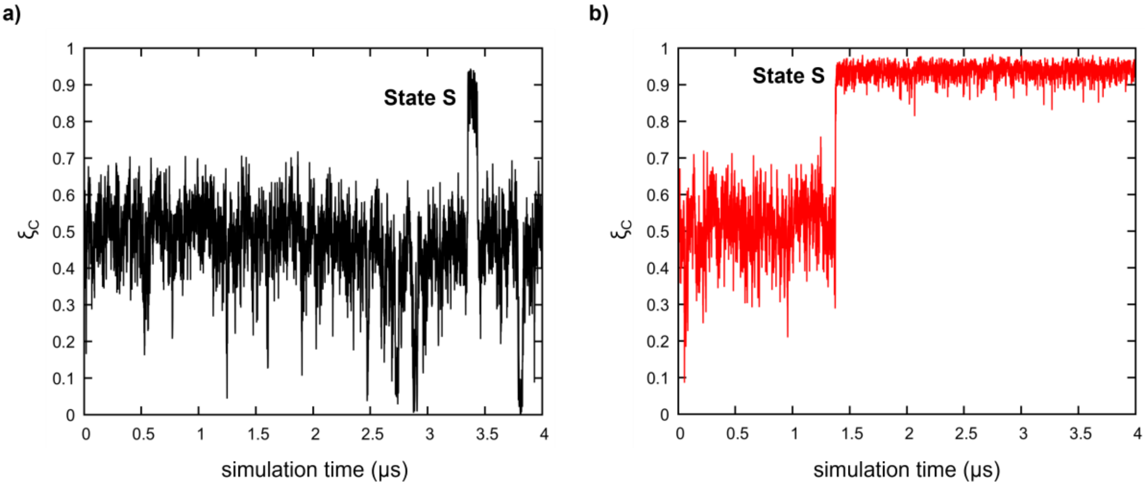


**Figure S17.** The collective variable $\xi_C$ as a function of simulation time obtained from unbiased MD simulations. a) A representative example of the $\xi_C$ evolution in the **S1** system (no cholestorol): the stalk forms after a few μs, but has a very short lifetime. b) A representative example of the $\xi_C$ evolution in the **S2** system (30% of cholesterol): once the stalk is formed, it remains stable for the rest of the simulation.
